# Trait repetitive negative thinking in depression is associated with functional connectivity in negative thinking state rather than resting state

**DOI:** 10.1101/2023.03.23.533932

**Authors:** Masaya Misaki, Aki Tsuchiyagaito, Salvador M. Guinjoan, Michael L. Rohan, Martin P. Paulus

## Abstract

Resting-state functional connectivity (RSFC) has been proposed as a potential indicator of repetitive negative thinking (RNT) in depression. However, identifying the specific functional process associated with RSFC alterations is challenging, and it remains unclear whether alterations in RSFC for depressed individuals are directly related to the RNT process or to individual characteristics distinct from the negative thinking process per se. To investigate the relationship between RSFC alterations and the RNT process in individuals with major depressive disorder (MDD), we compared RSFC with functional connectivity during an induced negative-thinking state (NTFC) in terms of their predictability of RNT traits and associated whole-brain connectivity patterns using connectome-based predictive modeling (CPM) and connectome-wide association (CWA) analyses. Thirty-six MDD participants and twenty-six healthy control participants underwent both resting state and induced negative thinking state fMRI scans. Both RSFC and NTFC distinguished between healthy and depressed individuals with CPM. However, trait RNT in depressed individuals, as measured by the Ruminative Responses Scale-Brooding subscale, was only predictable from NTFC, not from RSFC. CWA analysis revealed that negative thinking in depression was associated with higher functional connectivity between the default mode and executive control regions, which was not observed in RSFC. These findings suggest that RNT in depression involves an active mental process encompassing multiple brain regions across functional networks, which is not represented in the resting state. Although RSFC indicates brain functional alterations in MDD, they may not directly reflect the negative thinking process.

## Introduction

Repetitive negative thinking (RNT) is defined as a perseverative thought process that focuses on one’s problems or negative experiences in the present, past, and future (Ehring, 2021; Ehring and Watkins, 2008). Rumination and worry are the two main clinical types of RNT, with rumination referring to a passive, repetitive and evaluative focus on the symptoms of distress, whereas worry is a chain of thoughts focused on possible future negative outcomes (Nolen-Hoeksema et al., 2008). Rumination is often seen in individuals with Major Depressive Disorder (MDD) and has a moderate genetic correlation with MDD (Johnson et al., 2014). RNT predicts the onset of new episodes and the maintenance of existing symptoms of depression and is associated with reduced treatment response (Nolen-Hoeksema et al., 2008; Watkins and Roberts, 2020). The increased morbidity and treatment resistance of MDD individuals with RNT create a forceful impetus to identify the neurobiological underpinnings of RNT.

The neural correlates of RNT have been investigated using fMRI resting-state functional connectivity (RSFC), with the assumption that the resting state may indicate intrinsic brain functional alterationss in depression (Hamilton et al., 2015; Zhang et al., 2021). Given that RNT involves the process of perseverative thinking of self-referential negative thoughts (Ehring, 2021), many studies have focused on the default mode network (DMN) areas that have been implicated in self-referential thinking (Hamilton et al., 2015). For example, increased RSFC in DMN regions has been frequently reported to be associated with high RNT in depression (Bessette et al., 2018; Hamilton et al., 2015; Jacob et al., 2020; Makovac et al., 2020; Misaki et al., 2020; Stern et al., 2022; Wise et al., 2017; Yang et al., 2022; Zhu et al., 2017). RNT has also been associated with altered RSFC between the DMN and other regions, including the dorsolateral prefrontal cortex (Ichikawa et al., 2020; Peters et al., 2016), the amygdala (Connolly et al., 2013; Peters et al., 2016; Satyshur et al., 2018), and other executive control network nodes (Connolly et al., 2013; Feurer et al., 2021; Satyshur et al., 2018).

While these findings suggest that RSFC reflects brain functional alterations associated with rumination in depression, it is inherently difficult to identify the specific functional process associated with RSFC alterations. It remains unclear whether the RSFC associated with rumination in depression is directly related to the RNT process or to individual characteristics distinct from the negative thinking process itself. Resting-state fMRI does not necessarily reflect the true resting state of the brain, but rather a state that is not constrained by a specific task (Finn, 2021; Gonzalez-Castillo et al., 2021) and can vary over time (Greene et al., 2018; Lurie et al., 2020). Thus, RSFC can be an unstable and functionally unconstrained measure as an indicator of individual traits. Indeed, a study with large datasets including the ABCD study, Huma Connectome Project (HCP), and UK Biobank showed that the reproducibility of RSFC associations with individual traits could be low with a small sample size in a mass-univariate analysis (Marek et al., 2022). Meta- and mega-analysis studies of large cohort data of participants with MDD have also reported inconsistent RSFC alterations in the depressed group compared to previous studies (Goldstein-Piekarski et al., 2022; Tozzi et al., 2021; Yan et al., 2019; Zhang et al., 2020). These studies indicated either decreased or no difference in DMN RSFC in depressed individuals, while previous reports showed increased DMN RSFC in depression (for review: (Kaiser et al., 2015; Mulders et al., 2015; Williams, 2016)).

Several studies have also suggested that task-active states provide more informative insights into the neurophysiological associations of individual cognitive traits compared to the resting state (Greene et al., 2020; McCormick et al., 2022). For example, Finn and Bandettini (2021) demonstrated that FC patterns during movie-watching were more predictive of individual cognitive scores, specifically the principal component of the scores in the cognitive domain test, using data from the HCP. Similarly, Greene et al. (2018) found that FC during task-active states outperformed RSFC in predicting individual fluid intelligence scores. Considering these findings, it is plausible to suggest that brain functional changes associated with RNT in depression may exhibit more prominent effects during task-active states compared to the resting state. Indeed, task-based studies using a rumination-inducing task have also found differences in brain activity between individuals with high and low levels of RNT in depression. For example, a study focusing on the DMN found decreased connectivity within DMN subsystems during rumination (Chen et al., 2020). Functional changes related to RNT in a task-active state have also been observed in regions other than the DMN. These include connectivity between the salience and anterior parietal networks, where low connectivity between these networks was associated with high RNT following a sad event (Lydon-Staley et al., 2019). Additionally, reward-related regions showed a positive correlation between ventral striatum response and rumination (Erdman et al., 2020; Jones et al., 2017). Furthermore, increased connectivity between the angular gyrus and the rostrolateral prefrontal cortex region was associated with high rumination in the DMN and executive control regions (Jones et al., 2017). These findings suggest that the rumination induction task engages not only the DMN, but also regions of the salience and executive control networks that are not active during the resting state.

Although altered brain function associated with RNT in depression has been studied both at rest and during negative thinking (NT), it remains unclear which brain state better characterizes clinical RNT, and whether findings in these states reflect the same or independent functional changes. One could argue that functional changes during an NT task may better elucidate the pathology of depression because RNT is characterized by a negative thinking style (Nolen-Hoeksema et al., 2008). However, asking participants to engage in negative thinking may reduce the differences in brain states between individuals with and without RNT, and the induced negative thinking may not reflect the trait RNT. As such, the sensitivity for detecting RNT-related changes in brain function may be lower in the NT state than in the resting state. In fact, previous resting-state studies have often assumed that RSFC changes in MDD may reflect frequent spontaneous rumination at rest (Bessette et al., 2018; Connolly et al., 2013; Feurer et al., 2021; Hamilton et al., 2015; Jacob et al., 2020; Lois and Wessa, 2016; Nejad et al., 2013; Satyshur et al., 2018; Zhang et al., 2020; Zhang et al., 2022; Zhu et al., 2017). If this assumption is true, then resting and NT states should show similar changes in FC in depressed individuals, and FC in both states may be indicative of the severity of RNT in depression. However, if the resting and NT states reflect separate functional changes in depression, we should consider the implications of each state separately and which state better reflects the clinical trait of RNT.

The present study aims to investigate whether the resting state or the NT state is more associated with trait RNT in depression, and whether their changes reflect the same or different pathology. To achieve these goals, we utilized connectome-based predictive modeling (CPM) (Shen et al., 2017) and connectome-wide association (CWA) analysis (Shehzad et al., 2014). CPM is a machine learning approach used to create a predictive model of the brain-behavior relationship from whole-brain FC patterns. Multivariate predictive modeling approaches like CPM can overcome the small effect size problem of brain-behavior associations in mass-univariate analyses (Marek et al., 2022; Rosenberg and Finn, 2022) because they can aggregate univariate features to improve both sensitivity and robustness (Finn and Rosenberg, 2021; Rosenberg and Finn, 2022; Taxali et al., 2021). We applied CPM to whole-brain functional connectivity patterns in the resting and NT states to (1) discriminate individuals with MDD from healthy controls (HC) and (2) predict individual differences in trait RNT score, as measured by the Ruminative Response Scale (RRS) (Nolen-Hoeksema and Morrow, 1991). The RRS can be divided into three subscales: depressive, brooding, and reflective (Treynor et al., 2003). Because the brooding subscale most closely reflects the trait RNT, we focused on this subscale (RRS-B) in the present analysis. By comparing CPM performance between the resting and NT states, we could examine which state was more informative for characterizing depression and trait RNT.

Furthermore, we employed a connectome-wide association (CWA) approach (Shehzad et al., 2014) using longitudinal multivariate distance matrix regression (MDMR) analysis (Misaki et al., 2018) to comprehensively investigate FC changes in depression during both resting and NT states. MDMR (Anderson, 2001) is a multivariate analysis that examines associations between brain activity patterns and behavior across the entire brain. Like the CPM approach, CWA is a multivariate method that overcomes the limitations of mass-univariate analyses and provides a valuable complement to the CPM results. While CPM selects FCs that are sufficient for improved classification and prediction, potentially missing the comprehensive FC patterns associated with depression and RNT, CWA with MDMR quantifies the entire connectivity associated with depression and RNT in the voxel-to-voxel connectivity patterns of the whole brain. By combining these approaches, we evaluated the functional implications of FC changes in MDD during both resting and NT states, as well as their association with individual RNT traits.

## METHODS

### Participants

Twenty-eight medically and psychiatrically healthy (HC) individuals and forty-two individuals with MDD participated in the study. Each participant performed resting and rumination-inducing negative thinking (NT) tasks, which followed by other task runs including neurofeedback training (Tsuchiyagaito et al., 2023; Tsuchiyagaito et al., 2021). The present study analyzed their fMRI data during the resting-state and NT tasks. MDD participants met the DSM-5 (American Psychiatric Association, 2013) criteria for unipolar MDD based on the Mini-International Neuropsychiatric Interview 7.0.2 (Sheehan et al., 1998), and had current depressive symptoms with Montgomery-Åsberg Depression Rating Scale (MADRS) score > 6 (Montgomery and Åsberg, 1979). See Tsuchiyagaito et al. (2021) and Tsuchiyagaito et al. (2023) for detailed inclusion and exclusion criteria. The study protocols were approved by the Western Institutional Review Board, and all participants gave informed consent to participate in the study.

Two HC and six MDD participants were excluded from the analysis due to excessive head motion (more than 30% of time points (TR) censored with > 0.2 mm frame-wise displacement threshold in image processing) in either the resting or NT task. As a result, data from 26 HC participants (20 females, mean age = 23 years) and 36 MDD participants (28 females, mean age = 34 years) were included in the analysis. Supplemental Material Table S1 presents participant demographics and rumination and depression symptom scores.

### Scanning sessions and imaging parameters

The scanning session began with an anatomical scan, followed by a 6m50s resting state session and a 6m50s rumination-inducing NT task. In the resting state session, participants were instructed to clear their minds and not think about anything in particular while looking at the cross sign on the screen. In the NT task, participants were instructed to think of a recent time when they felt rejected by someone who meant a lot to them while looking at the cross sign on the screen. The instruction provided for the NT task aimed to elicit a typical rumination process by focusing on common triggers of rumination such as personal relationships, past mistakes, negative experiences, and social interactions (Joubert et al., 2022). Participants completed these scans before any other task sessions so that no effect of other tasks confounded the resting state and NT sessions. Also, the resting state session always preceded the NT task so that no effect of the NT task was confounded with the resting state scan.

MRI scans were performed using a GE 3 Tesla MR750 Discovery scanner (GE Healthcare, Milwaukee WI, USA). The anatomical scan acquired a T1-weighted image with the MPRAGE sequence of TR/TE = 5/2 ms, SENSE acceleration R = 2, flip angle = 8, delay/inversion time TD/TI = 1400/725 ms, sampling bandwidth = 31.2 kHz, FOV = 240 x 192 mm, 124 axial slices, slice thickness = 1.2 mm, and scan time = 4 min 59 s. The resting and NT session functional scans acquired T2*-weighted images with a gradient echo planar sequence of TR/TE = 2000/25 ms, flip angle = 90, SENSE acceleration R = 2, acquisition matrix = 96 x 96, FOV/slice = 240/2.9 mm, and scan time = 6m50s (205 TRs).

### MRI image processing

Analysis of Functional NeuroImages package (AFNI; http://afni.nimh.nih.gov/) (Cox, 1996) was used for MRI image processing. The same processing pipeline was used for both resting and NT state data. The first three volumes of functional images were excluded from the analysis. Processing included despiking, RETROICOR (Glover et al., 2000) and respiratory volume per time (Birn et al., 2008) physiological noise corrections, slice timing alignment, motion alignment, nonlinear warping to the MNI template brain with resampling to 2mm^3^ voxel volume using the ANTs (https://picsl.upenn.edu/software/ants/) (Avants et al., 2008), spatial smoothing with a 6mm-FWHM Gaussian kernel within the brain mask, and scaling of the signal to the percent change relative to the mean in each voxel. General linear model (GLM) analysis was then applied with censoring volumes with > 0.2mm frame-wise displacement and regressors of Legendre polynomial models of slow signal fluctuations, 12 motion parameters (3 shifts, 3 rotations, and their temporal derivatives), three principal components of ventricular signals, and local white matter average signal (ANATICOR) (Jo et al., 2010). The residual of the GLM analysis was used as the processed fMRI data for the calculation of functional connectivity.

### Brain parcellation for functional connectivity matrix calculation

The Shen 268-node atlas (Finn et al., 2015; Shen et al., 2013) was used to parcellate brain regions. We excluded 38 regions around the orbitofrontal, ventricles, and in the lower part of the brain from the analysis, because they were not covered by functional images or showed significant signal loss in many participants. The map of the excluded regions and their region indices in the Shen 268-node atlas are shown in Supplemental Material Figure S1.

Functional connectivity between the 230 regions was calculated for the mean signals in the regions using Pearson’s correlation followed by Fisher’s z-transformation. The upper triangular part of the connectivity matrix, 26335 values, was the input for the connectome-based predictive modeling (CPM) analysis.

### Connectome-based Predictive Modeling (CPM)

CPM analysis was conducted for two distinct tasks: the classification of individuals with MDD and HC, and the prediction of RRS-B scores. Separate models were constructed for each task. The effects of age, sex, and average motion (frame-wise displacement) were regressed and eliminated from the connectivity values in both resting-state functional connectivity (RSFC) and negative thinking functional connectivity (NTFC) data. These covariate effects were also removed from the RRS-B scores in the RRS-B prediction task.

For the MDD-HC classification task, CPM constructed a prediction model using the following steps: 1) selecting connectivity values with a large absolute t-value for the difference between the MDD and HC groups (the *t*-value threshold was optimized using nested cross-validation), 2) summing the selected connectivity values for each positive and negative *t*-value in each participant, and 3) fitting a logistic regression model to classify MDD and HC individuals based on the summed scores of positive and negative connectivities, respectively (Shen et al., 2017). The performance of the classification model was assessed using the Area Under the Curve (AUC) of the receiver operating characteristic (ROC) curve.

For the RRS-B prediction task, the same procedure was employed to construct a prediction model as in the classification task, except that connectivity values with a high absolute Pearson’s correlation (*r*) with the RRS-B scores were selected at step 2). Then, a linear regression model was fitted to predict the RRS-B scores at step 3), following the same procedure described above. The performance of the RRS-B prediction was evaluated using Spearman’s correlation coefficient to measure the association between the true and predicted RRS-B scores. The RRS-B prediction was performed only for the MDD group because prediction for all participants, including both MDD and HC, could be confounded by significant group differences in RRS-B scores.

The classification and prediction performances were assessed using 5-fold cross-validation, where participants were randomly divided into training (80%) and test (20%) sets. The model hyperparameters, such as the absolute *t*-value and absolute correlation thresholds for connectivity selection, were optimized through a nested 4-fold cross-validation within the training set. The training set was further divided into a training subset (75%) and a validation subset (25%) for this purpose. Grid search was employed to explore different values for the hyperparameters: *t*-values ranging from 1.0 to 5.0 with intervals of 0.5 for MDD-HC classification, and correlation values (*r*) ranging from 0.1 to 0.5 with intervals of 0.05 for RRS-B prediction. The entire process of 5-fold cross-validation was repeated 100 times to obtain a reliable estimate of predictive performance.

A permutation test was performed to assess the statistical significance of the results. The output values were randomly permuted 1000 times, and in each permutation iteration, 5-fold cross-validation was repeated 20 times with different random splits. The same hyperparameter optimization procedure with nested cross-validation was also applied during the permutation test. The median of 20 repeats was obtained in each iteration to create a null distribution.

### Connectome-wide association analysis

For a comprehensive functional connectivity investigation of the differences between rest and NT states, between HC and MDD, and the RRS-B association, we performed multivariate distance matrix regression (MDMR) analysis on whole-brain voxel-to-voxel connectivity (Shehzad et al., 2014). MDMR is a variant of MANOVA that uses nonparametric statistics (Anderson, 2001). The analysis computes connectivity maps for a seed voxel, and the distance matrix of the whole-brain connectivity maps across samples (i.e., participant x run) is used as the multivariate dependent variable for the linear model with multiple explanatory factors. These procedures are repeated for each voxel as a seed, and the regression statistic is assigned to the seed voxel.

The processed fMRI images were resampled into 4mm^3^ voxels, and the seed and its connectivity map were constrained in the gray matter voxels. The distance matrix between the resting-state and NT-state connectivity maps for all participants, calculated by Euclid distance, was the dependent variable in the MDMR. We used the longitudinal design introduced by Misaki et al. (2018) to account for the within-subject factor of resting and NT runs. The model included state (rest, NT), group (MDD, HC), RRS-B score, interactions of these three factors (including all two-ways and three-way), age, sex, motion size (mean FD), and subject-specific factor variables (Misaki et al., 2018; Winkler et al., 2014). The significance of the MDMR statistic of the pseudo-*F* value (Anderson, 2001) was assessed by permutation test with 10,000 repeats. The map was thresholded by voxel-wise *p* < 0.001 and family-wise error correction by cluster-extent *p* < 0.05. The cluster-extent threshold was also evaluated by permutation test.

The regions showing a significant effect in the sum of the factors of interest (state [resting, NT], group [MDD, HC], RRS-B, and their interaction) were selected for post-hoc seed-based connectivity analysis. We opted to calculate the sum of the effects of interest in our approach because analyzing individual factors with interaction terms and evaluating pseudo *F*-values and *p*-values for each of them would require a complex and time-consuming process (Fox, 2015; Langsrud, 2003). Instead, the effect of each factor was delineated in the post-hoc seed-based analysis using a linear mixed-effect model.

The post-hoc analysis was performed in the original image resolution. Seed regions were placed at peak locations of the significant cluster in the MDMR statistical map for the sum of the effects of interest by extracting peak coordinates separated by at least 30mm in a cluster. The seed region was defined by a sphere of 6mm radius centered on the peak coordinates. The mean signal time course of the seed region was used as a reference signal to calculate connectivity with other voxels in the whole brain. For the statistical testing of the post hoc analysis, we used linear mixed effect (LME) model analysis with the same linear model specification as the MDMR, except that the subject factor was entered as a random effect on the intercept. We used the lme4 package (Bates et al., 2015) with the lmerTest package (Kuznetsova et al., 2017) in the R language and statistical computing (R Core Team, 2022) for the LME analysis.

We note that the post-hoc analysis was performed to elucidate the connectivity map associated with the MDMR result, and the MDMR is the test for the global connectivity pattern, not for individual voxel-wise connectivity. Therefore, we used a lenient voxel-wise threshold (*p* < 0.05) in the post-hoc evaluation to illustrate the global connectivity pattern for the seed with significant MDMR statistics. Nevertheless, we evaluated the cluster size threshold with this voxel-wise threshold using cluster size simulation with 3dClustSim in AFNI (Cox et al., 2017), so the result complies with the corrected *p* < 0.05 threshold. The map of each contrast (i.e., MDD-HC in each state, rest-NT in each group, and RRS-B association in each group and state) was calculated using the R emmeans package (Lenth, 2022).

## Results

### CPM prediction of HC and MDD groups

Figure 1A shows the distributions of AUC for the MDD-HC classification over 100 iterations of 5-fold cross-validation with different random splits. The median AUCs were 0.826 (*p* < 0.001) for the RSFC and 0.794 (*p* < 0.001) for the NTFC. The difference in performance between RSFC and NTFC was not significant (*p* = 0.686). Connectivity included in the prediction model is plotted on the glass brain (Fig. 1B) and with a circle plot (Fig. 1C). These plots show the connectivity selected more than 50% of the time in multiple (100 x 5) cross-validation iterations. The line color indicates the mean connectivity difference (z-value) between MDD and HC groups (warm color indicates higher connectivity for MDD). In Figure 1C, the network labels are adapted from Drysdale et al. (2017). Note that these maps are presented only to illustrate the FC patterns used by CPM as a whole, and not to evaluate the individual FC association with the classification. As CPM is a multivariate pattern analysis, it is not appropriate to evaluate individual FC associations independently. Therefore, we did not conduct a strict statistical test to evaluate the independent effect of each FC.

**Figure 1.**
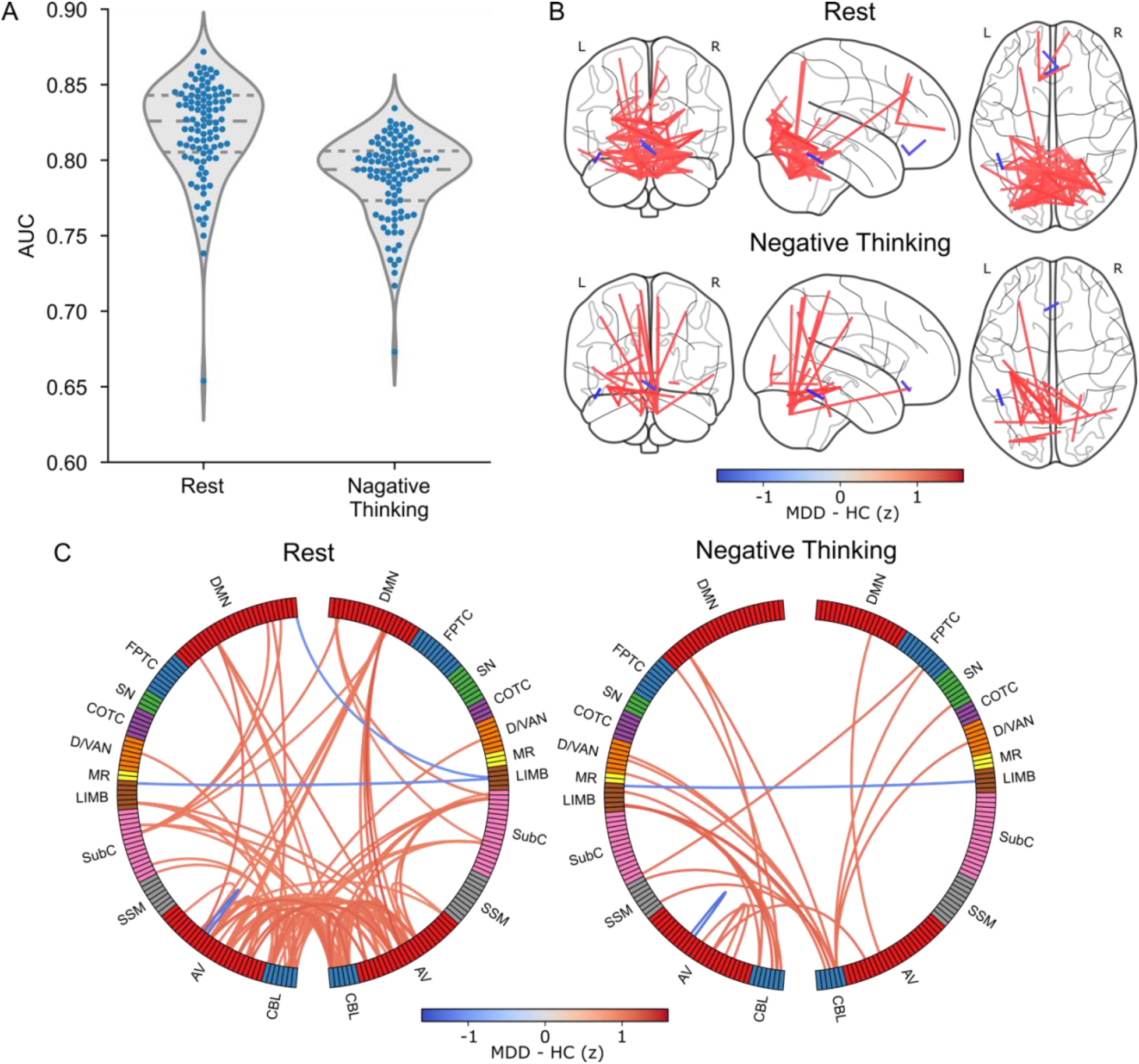
A. CPM prediction performance distributions for MDD-HC classification. Each point indicates one iteration of the 5-fold cross-validation result (100 iterations with different random splits were performed). The violin plot and horizontal lines indicate the distribution curve and quartile positions, respectively. B. Connectivity selected by the CPM model. The connectivities selected by more than 50% cross-validation iterations were plotted on the glass brain. The line color indicates the connectivity difference (z-value) between the MDD and HC groups. C. Circle plots of the same connectivities as in B summarized for each network region. Network labels are adapted from (Drysdale et al., 2017). DMN: Default Mode Network, FPTC: Fronto-Parietal Task Control, SN: Salience Network, COTC: Cingulo-Opperculum Task Control, D/VAN: Doral Visual Attention Network, MR: Memory Retrieval, LIMB: default mode/limbic, SubC: Subcortical, SSM: Sensory SomatoMotor, AV: Auditory-Visual, CBL: Cerebellum.

In the resting state, CPM classified participants as MDD based on high connectivity within visual cortex and cerebellar regions and their connectivity to DMN regions. In the circle plot (Fig. 1C), the one cool-colored line (represented by the bilateral LIMB [default mode/limbic] connection) indicated the reduced functional connectivity between the bilateral subgenual anterior cingulate cortex (sgACC) regions in MDD compared to HC. In the NT state, the CPM used FCs in similar areas as in the resting state, while the number of FCs consistently selected across the cross-validation iterations was fewer than in the resting state (Figs. 1B and 1C).

### CPM prediction of individual RRS-B score

Figure 2A displays the distributions of Spearman’s correlations between true and predicted RRS-B scores for MDD over 100 iterations of 5-fold cross-validation with different random splits (Supplemental Material Fig. S2 shows CPM prediction results including both groups). The median Spearman’s correlations were -0.049 (*p* = 0.447) for RSFC and 0.279 (*p* = 0.041) for NTFC. The difference in performance between the resting and NT states was significant (*p* = 0.043). Figures 2B and 2C show the connectivity included in the prediction model that was selected more than 85% of the time in the cross-validation iterations (plots with 50% threshold are shown in Supplemental Material Fig. S3). Line color indicates FC correlation with RRS-B (z-transformed, warm color indicates positive correlation).

**Figure 2.**
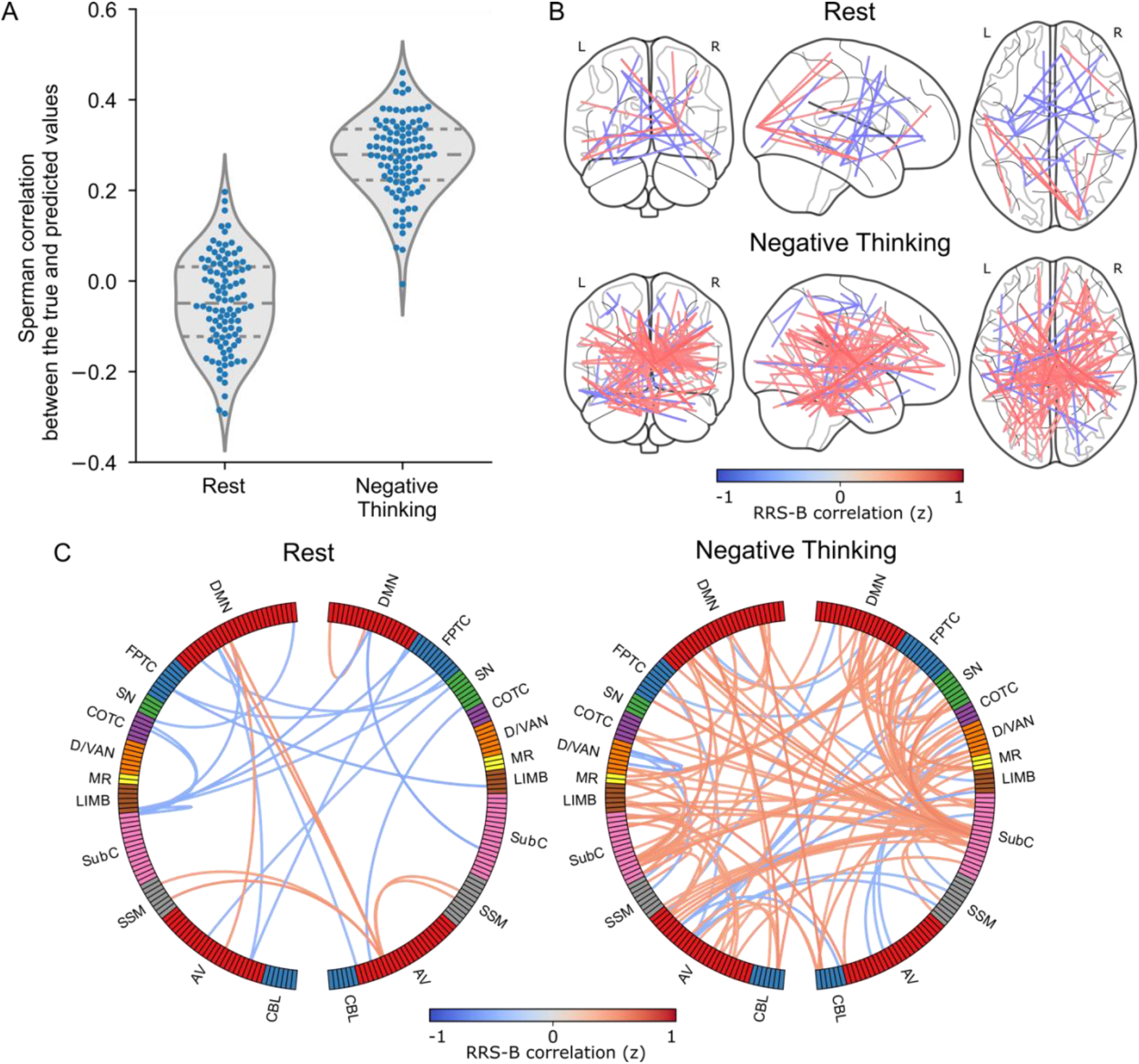
A. Distributions of the CPM prediction performance for individual RRS-B scores in the MDD group. Each point indicates one iteration of the 5-fold cross-validation result (100 iterations with different random splits were performed). The violin plot and horizontal lines indicate the distribution curve and quartile positions. B. Plots of the connectivity selected by the CPM model in more than 85% of the cross-validation iterations. Line color indicates the connectivity correlation with RRS-B (z-transformed). C. Circle plots of the same connectivities as in B, summarized for each network region. Network labels are adapted from Drysdale et al. (2017) (see Fig. 1 for the network label abbreviations).

In the NT state, connectivity associated with RRS-B prediction was distributed over broad brain areas with high consistency across participants (across cross-validation). The higher connectivity from the thalamus regions (SubC nodes with dense, warm lines in Fig. 2C) in the NT state characterizes high RRS-B individuals in the MDD group. In contrast, in the resting state, many connectivities consistently selected by CPM were negatively correlated with RRS-B (Fig. 2B and Supplemental Material Fig. S3B).

### MDMR connectome-wide association analysis

Figure 3 shows the regions with significant effects of interest (sum of the effects of state, group, RRS-B, and their interactions) with voxel-wise *p* < 0.001 and cluster-extent corrected *p* < 0.05 in the MDMR analysis. Significant clusters were found in DMN regions (i.e., precuneus, posterior cingulate cortex [PCC], medial prefrontal cortex [MPFC]), executive control regions (i.e., supplementary motor area [SMA] and lateral frontal regions including inferior frontal gyrus [IFG]), the caudate region, and the cerebellum. Post-hoc seed-based connectivity analysis was performed on the peak areas in these significant clusters. Table 1 shows the seed points used for the post-hoc analysis.

**Figure 3.**
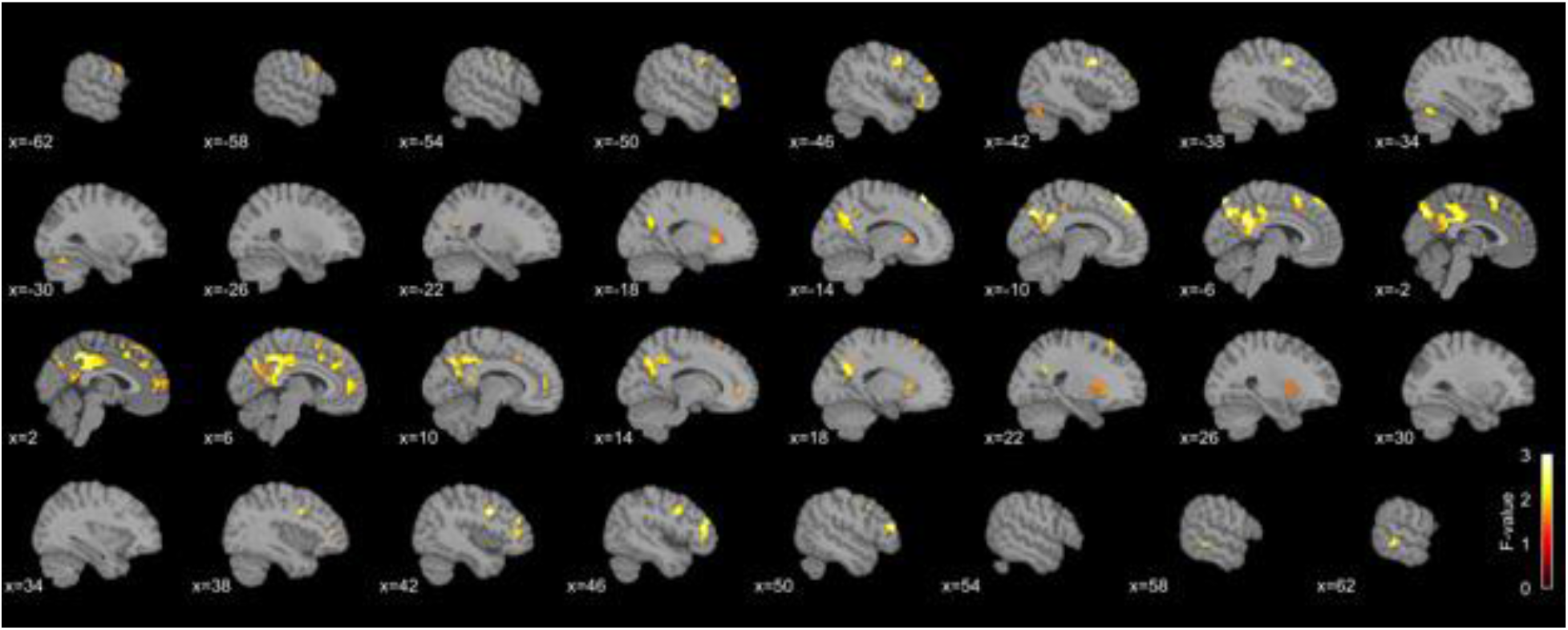
Significant regions with the MDMR statistics for the sum of the effects of interest (state, group, RRS-B, and their interactions) with voxel-wise *p* < 0.001 and cluster-extent corrected *p* < 0.05.

**Table 1.**
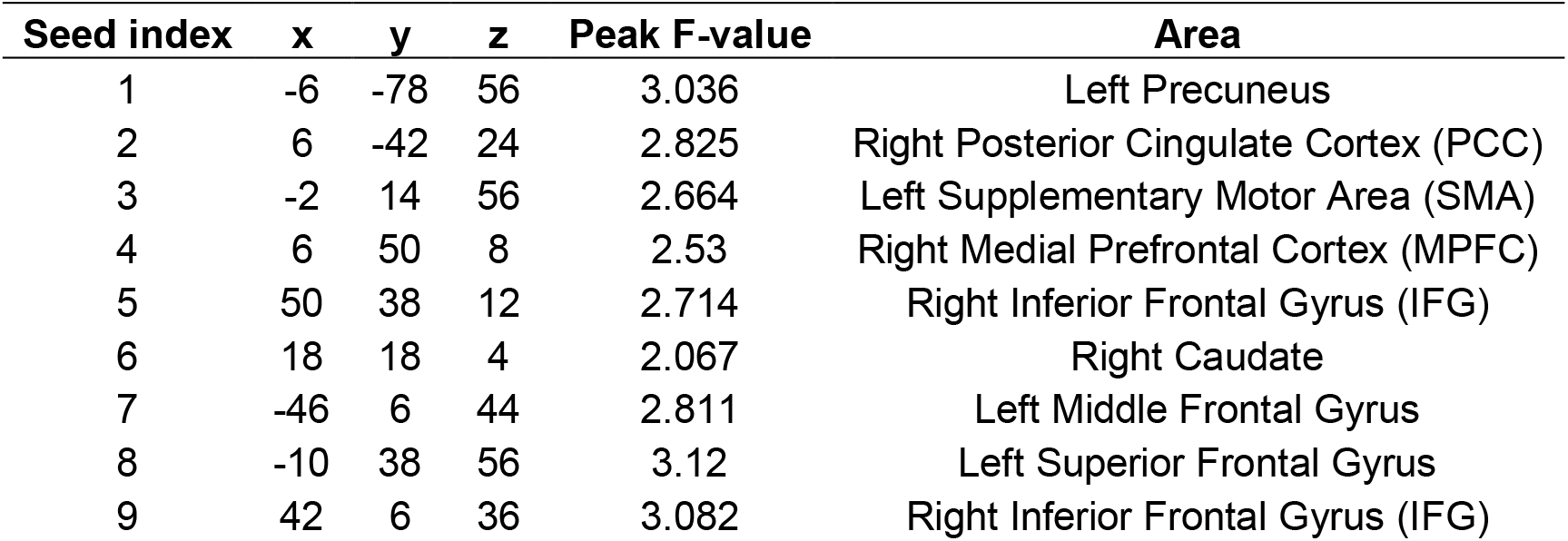
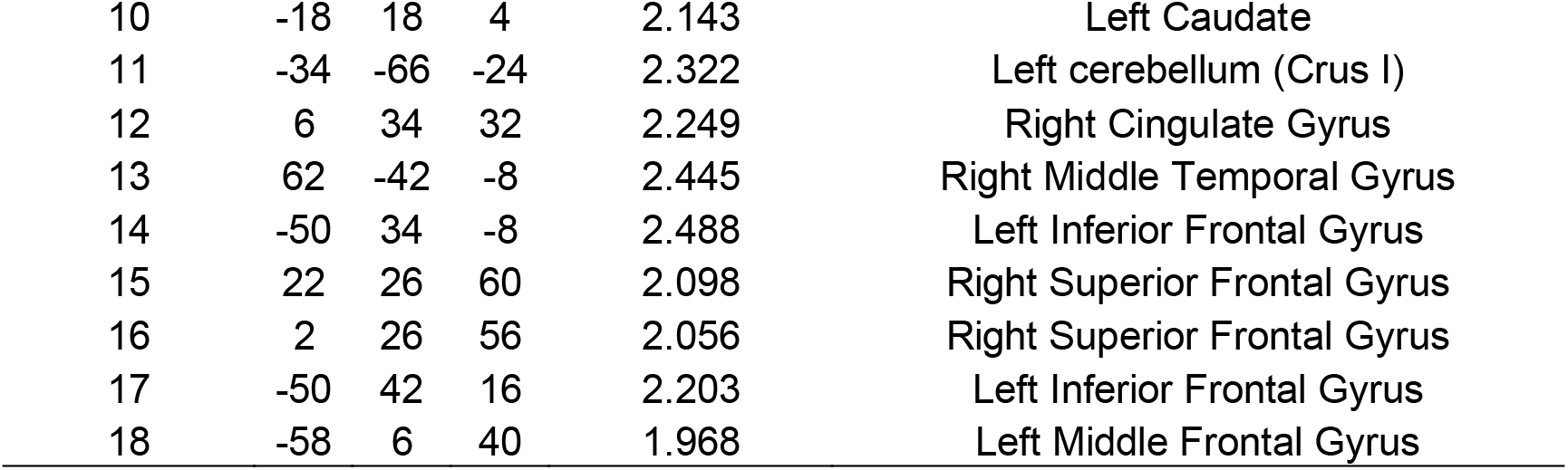
The seed points in the significant clusters of the MDMR statistics.

| Seed index | x | y | z | Peak F-value | Area |
| --- | --- | --- | --- | --- | --- |
| 1 | -6 | -78 | 56 | 3.036 | Left Precuneus |
| 2 | 6 | -42 | 24 | 2.825 | Right Posterior Cingulate Cortex (PCC) |
| 3 | -2 | 14 | 56 | 2.664 | Left Supplementary Motor Area (SMA) |
| 4 | 6 | 50 | 8 | 2.53 | Right Medial Prefrontal Cortex (MPFC) |
| 5 | 50 | 38 | 12 | 2.714 | Right Inferior Frontal Gyrus (IFG) |
| 6 | 18 | 18 | 4 | 2.067 | Right Caudate |
| 7 | -46 | 6 | 44 | 2.811 | Left Middle Frontal Gyrus |
| 8 | -10 | 38 | 56 | 3.12 | Left Superior Frontal Gyrus |
| 9 | 42 | 6 | 36 | 3.082 | Right Inferior Frontal Gyrus (IFG) |
| 10 | -18 | 18 | 4 | 2.143 | Left Caudate |
| 11 | -34 | -66 | -24 | 2.322 | Left cerebellum (Crus I) |
| 12 | 6 | 34 | 32 | 2.249 | Right Cingulate Gyrus |
| 13 | 62 | -42 | -8 | 2.445 | Right Middle Temporal Gyrus |
| 14 | -50 | 34 | -8 | 2.488 | Left Inferior Frontal Gyrus |
| 15 | 22 | 26 | 60 | 2.098 | Right Superior Frontal Gyrus |
| 16 | 2 | 26 | 56 | 2.056 | Right Superior Frontal Gyrus |
| 17 | -50 | 42 | 16 | 2.203 | Left Inferior Frontal Gyrus |
| 18 | -58 | 6 | 40 | 1.968 | Left Middle Frontal Gyrus |

Figure 4 summarizes the representative MDD-HC contrast in the post-hoc analysis for the two seeds at the DMN hub nodes; PCC (seed 2; seed index corresponds to Table 1) and MPFC (seed 4). Significant FC maps of all seeds are shown in Supplemental Material Figures S4 and S5 for the resting and NT states, respectively. In the resting state, MDD had higher connectivity than HC from these DMN seeds to the visual cortex regions (Figs. 4A and 4B). Other seeds also showed higher connectivity for MDD than HC in the occipital areas (Supplemental Material Fig. S4), which was consistent with the CPM results. In the NT state, higher connectivity was also seen in the occipital regions (Figs. 4C and 4D), although the lower connectivity areas for MDD than HC of these seeds seen in the resting state (cool color regions in Figs. 4A and 4B) were not seen in the NT state (Figs. 4C and 4D), indicating that the MDD-HC contrast decreased in the NT state compared to the resting state. Notably, PCC seed connectivity in the precuneus regions was higher for HC than MDD in the resting state (cool color regions in Figs. 4A), indicating that posterior DMN FC was higher for HC than MDD in the resting state. In addition, MDD group had higher connectivity between these DMN seeds and the IFG region in the NT state (Figs. 4C and 4D), which was not seen in the resting state.

**Figure 4.**
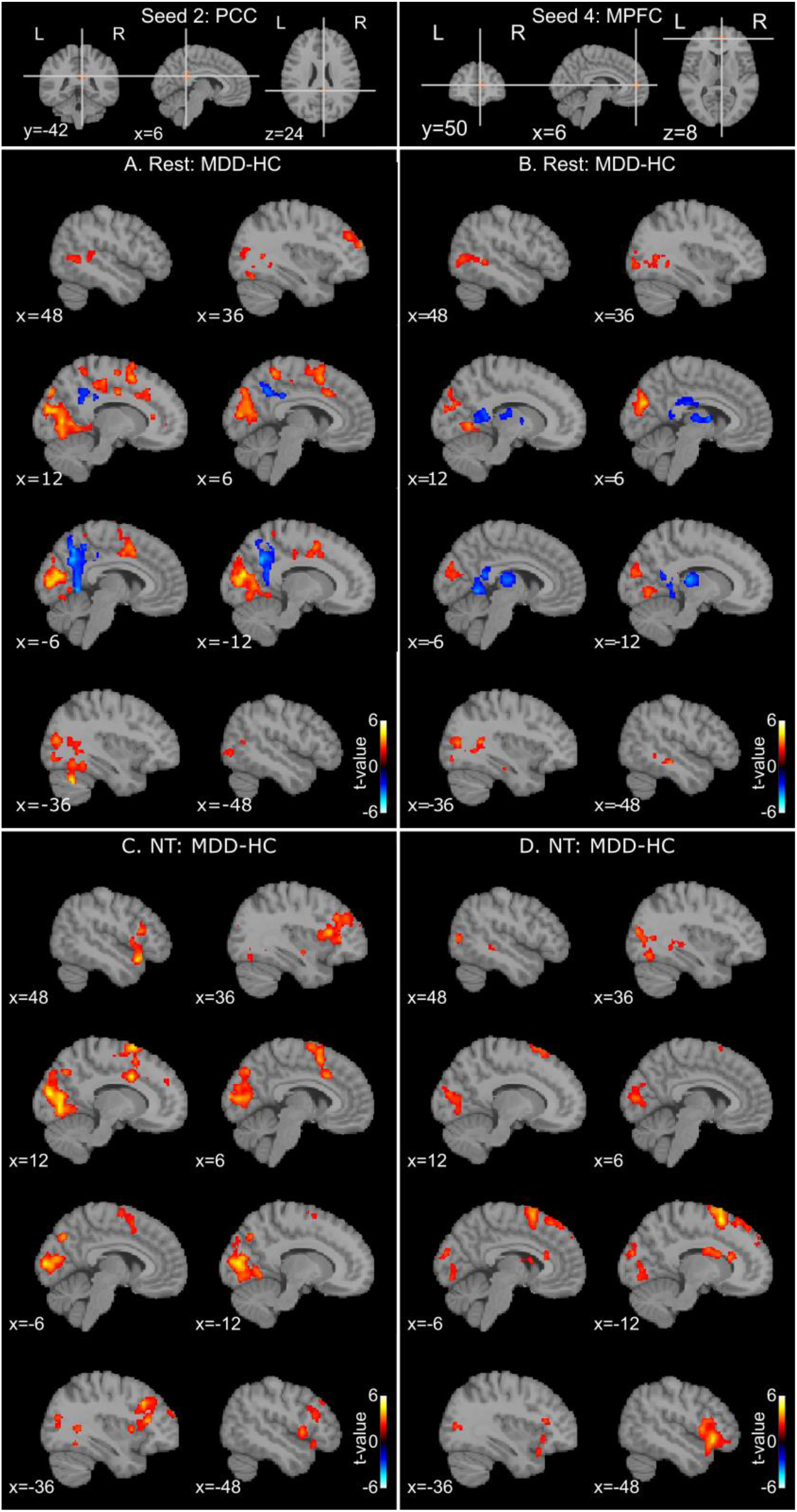
MDD-HC contrast connectivity maps in the MDMR post-hoc analysis with the PCC (seed 2) and MPFC (seed 4) seeds (top row) for the resting (A, B) and negative-thinking (NT; C, D) states. The map shows the *t*-value for the MDD-HC contrast. The seed index corresponds to Table 1. PCC: posterior cingulate cortex, MPFC: medial prefrontal cortex.

Figure 5 summarizes the representative NT-Rest contrast in the post-hoc analysis for three seeds, PCC (seed 2), SMA (seed 3), and right IFG (seed 9). Significant FC maps of all seeds are shown in Supplemental Material Figures S6 and S7 for HC and MDD, respectively. The PCC seed, a DMN hub region, had higher connectivity with regions in the executive control areas, including the lateral premotor and prefrontal regions, and the anterior insula in the NT state than in the resting state in MDD (Fig. 5D). The SMA and IFG seeds had higher connectivity with the precuneus area in the NT state than in the resting state in MDD (Figs. 5E and 5F). These indicate that connectivity between posterior DMN regions (precuneus and PCC) and executive control regions was increased in the NT state in MDD. This increased connectivity was not observed in HC (Figs. 5A, 5B and 5C). Connectivity between the SMA and precuneus showed an opposite pattern between HC and MDD (Figs. 5B and 5E); it was higher in rest than in NT for HC, but higher in NT than in rest for MDD. Connectivity within the posterior DMN regions was higher in the resting state than in the NT state in both HC and MDD (Figs. 5A and 5D).

**Figure 5.**
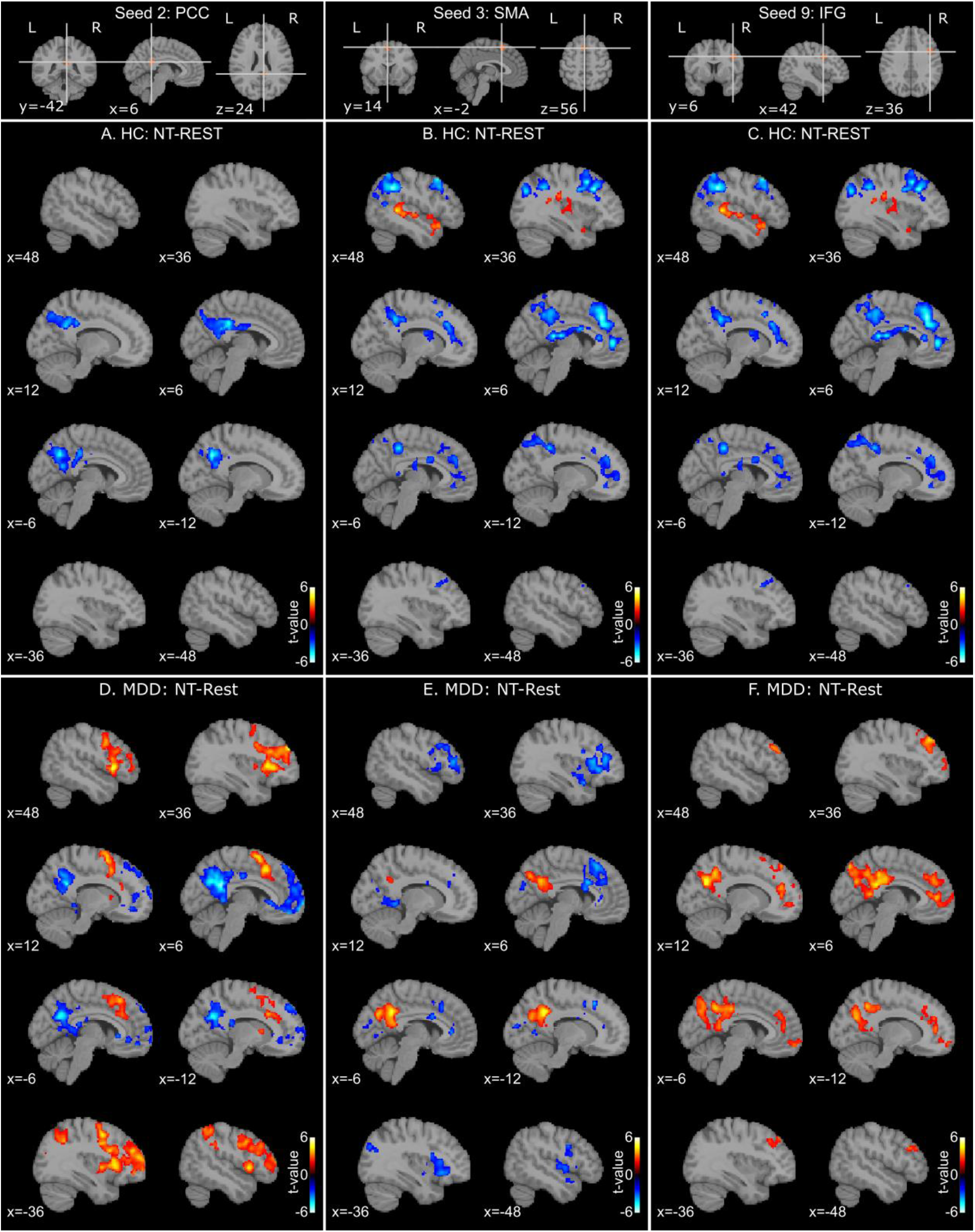
NT (negative thinking)-Rest contrast connectivity maps in the MDMR post-hoc analysis with the PCC (seed 2), SMA (seed 3), and right IFG (seed 9) seeds (top row) for the HC (A, B, C) and MDD (D, E, F) groups. The map shows the *t*-value for the NT-Rest contrast. The seed index corresponds to Table 1. PCC: posterior cingulate cortex, SMA: supplementary motor area, IFG: inferior frontal gyrus.

Figure 6 shows the RRS-B associations in the post hoc analysis for MDD. Significant FC maps of all seeds are shown in Supplemental Material Figs. S8 and S9 for resting and NT states, respectively. We did not examine the RRS-B association in HC because the score in HC did not have enough variance to assess the association robustly. The most striking observation was the connectivity of the cerebellum (seed 11). In the NT state, the connectivity of this cerebellar seed showed a positive correlation with the RRS-B in broad cortical areas, including DMN regions (i.e., PCC and MPFC, Fig, 6D). In contrast, in the resting state, cerebellar connectivity associated with the RRS-B was restricted to motor and premotor cortex and did not extend to the DMN (Fig. 6B). The negative RRS-B association with the FC between the precuneus and thalamus was also observed for the resting state in MDD (Fig 6A).

**Figure 6.**
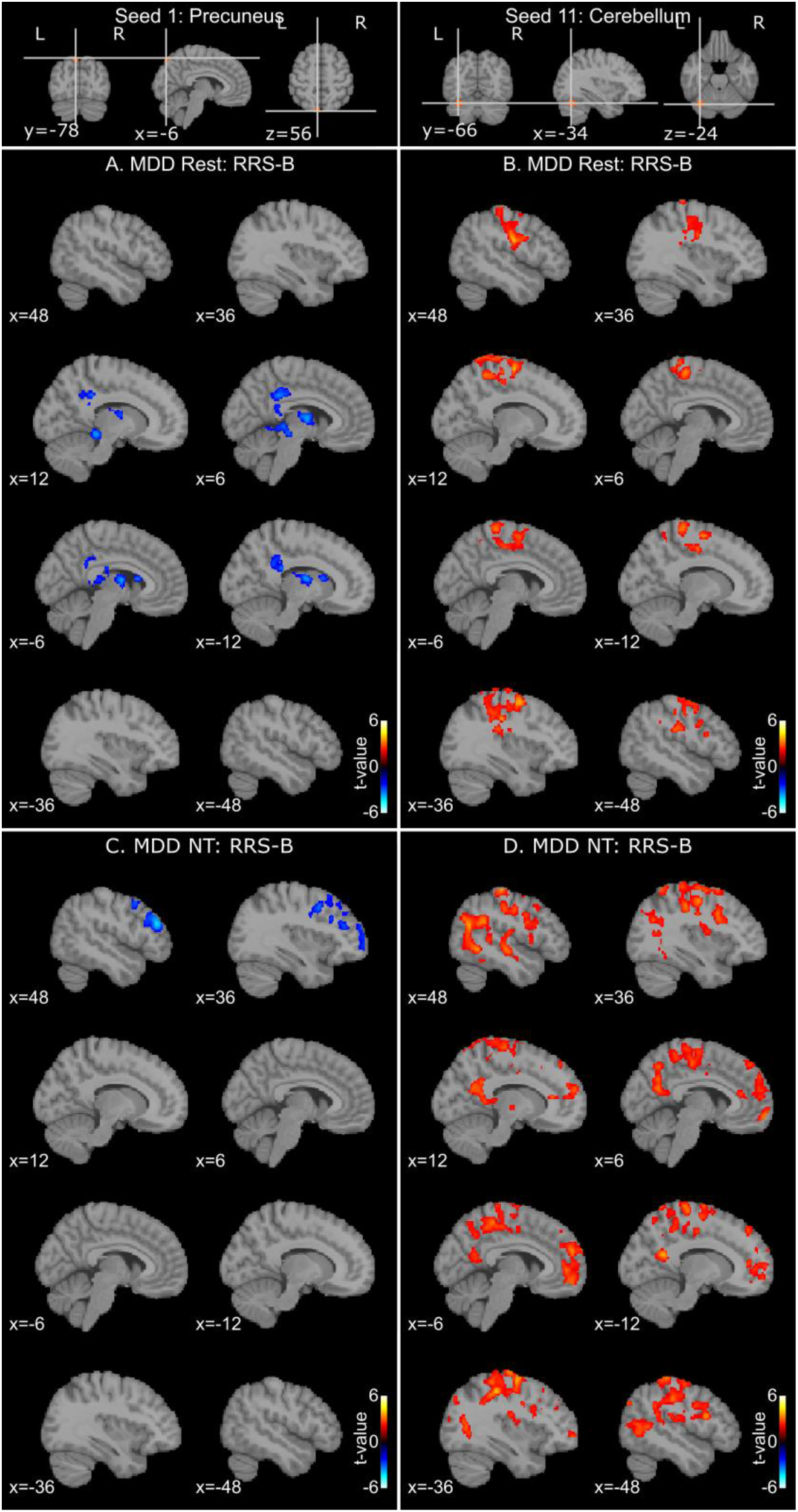
RRS-B association connectivity maps in the MDMR post-hoc analysis for seeds with the significant RRS-B association in the MDD group for the resting (A, B) and negative-thinking (NT; C, D) states. The map shows the *t*-value for the RRS-B association. The seed index corresponds to Table 1.

## Discussion

The primary findings of the present study are as follows: 1) CPM analysis demonstrated that both RSFC and NTFC were capable of distinguishing between HC and MDD individuals, 2) NTFC demonstrated predictive capability for trait RNT in individuals with depression, whereas RSFC did not show such predictive ability, and 3) CWA analysis indicated that the negative thinking process in MDD was associated with increased functional connectivity (FC) between regions of the default mode network and executive control regions, which was not observed in RSFC or in the HC group.

As both RSFC and NTFC were effective in differentiating individuals with MDD from HC, it is warranted to investigate functional brain alterations in MDD using resting-state measures. However, RSFC did not predict trait RNT in depression, suggesting that RSFC alterations in depression may not directly reflect the ongoing RNT process. This finding contradicts the assumption that modifications in RSFC in MDD arise from heightened spontaneous rumination during the resting state in these individuals. While RSFC indicates alterations in brain connectivity, further investigations are needed to understand the specific processes underlying the relationship between RSFC and RNT in depression.

Investigations of FC patterns associated with CPM prediction and complementary CWA analysis further revealed the significant difference between resting and NT states in depressed individuals. FCs distinguishing MDD from HC were found in areas of the visual cortex and cerebellum and their connections with DMN regions in both CPM and CWA analyses. Reduced FC between bilateral sgACC areas was also used by CPM to classify MDD. Altered sgACC function in depression has been reported for MDD (Brakowski et al., 2017; Drevets et al., 2008), and disrupted sgACC activity could reduce its bilateral functional coupling. While increased visual cortex activation during rumination has been reported in adolescents with remitted MDD (Burkhouse et al., 2017), a negative correlation between a trait rumination score (RRS) and visual cortex activation in both resting and task (face classification) conditions has also been reported (Piguet et al., 2014). Thus, while the increased FC in visual cortex in MDD may at least reflect higher visual imagery than HC at rest, it may not be specifically associated with negative thinking.

Interestingly, DMN connectivity was not selected in the CPM classification, and MDMR analysis revealed decreased posterior DMN connectivity in MDD compared to HC in the resting state. This is consistent with the meta- and mega-analysis studies of large cohort data that reported decreased or no difference in resting-state DMN FC in depressed individuals (Goldstein-Piekarski et al., 2022; Tozzi et al., 2021; Yan et al., 2019; Zhang et al., 2020).

In contrast to the MDD-HC classification, RRS-B prediction was well performed with NTFC, but not with RSFC. The FCs that were informative for predicting RRS-B in MDD in the CPM analysis were distributed over large areas of the brain (Supplemental Material Fig. S3). The involvement of many cortical regions, including the limbic and medial and dorsolateral prefrontal regions, has also been reported in the rumination induction task (Cooney et al., 2010). These suggest that RNT is associated with large-scale network and inter-network interactions (Lydon-Staley et al., 2019; Zhang et al., 2020) rather than a focal processing abnormality. MDMR results showing significant FC differences between resting and NT states in MDD support the perspective of multi-network involvement in the RNT process. Connectivity between the precuneus and executive control regions (i.e., SMA, IFG) and the salience network region (i.e., anterior insula) was increased in the NT state compared to the resting state in MDD participants (Fig. 5). Also, the FC between the DMN seeds and IFG in the NT state was significantly higher in MDD than in HC (Figs. 4C and 4D), suggesting that such an increase in the NT state is specific to depressed individuals.

When we used a strict threshold for plotting the FCs informative for predicting RRS-B, the dense connection was seen in the right thalamus area, which had a positive correlation with RRS-B (Fig. 2C). The significant RRS-B association with FC between thalamus and precuneus was also observed in the resting state in MDD (Fig. 6A), but in a negative direction, highlighting that the resting state in MDD had a significantly different FC pattern than the NT state. The involvement of the thalamus in RNT has been demonstrated in a 7T fMRI study (Steward et al., 2022), suggesting that the thalamus, with its extensive cortical pathways, may act to increase synchrony between cortical regions to maintain complex mental representations, including RNT. In addition, emerging clinical evidence suggests a role for right thalamic-cortical circuitry in the amelioration of depression in neuromodulation treatments (Lippitz et al., 1999; Riestra et al., 2011; Scangos et al., 2021), highlighting the potential clinical implications of this particular finding to help refine neuromodulatory procedures for MDD.

The RRS-B association in the CWA analysis was seen for the cerebellum seed connectivity with broad cortical regions. This cerebellar region (crus I) is functionally related to executive control network areas (Habas et al., 2009). A report of a blunted response of this region to reward anticipation in depressed individuals with high RNT (Park et al., 2022) also suggests that trait RNT may influence activation of this region. The associations of RRS-B with this seed to broad cortical regions, not limited to the executive control region but including the DMN regions in the NT state, suggest that the RNT is an active process involving multiple networks, not limited to the DMN. The increased FC between the SMA, a region that monitors and evaluates an active process (Bonini et al., 2014), and the precuneus, a region involved in self-referential thinking (Fig. 5E), also supports the idea that RNT in depression is an active process. The involvement of many cortical areas in the RNT process has also been reported in previous studies, including increased FC from the PCC to many cortical areas in the NT state compared to the resting state (Berman et al., 2014).

Discussing the limitations of the current study is warranted. The significant age difference between the MDD and HC groups may have biased the present results concerning the MDD-HC difference. To mitigate this, we excluded the age effect from the FCs in the CPM analysis, and age was included as a covariate factor in the MDMR analysis. However, excluding the age effect may have also removed the association between FC and depression if it interacted with age. Indeed, Andreescu et al. (2014) found an interaction between age and anxiety on DMN connectivity, where the effect of anxiety on FC was greater in older participants. Therefore, we acknowledge that the present findings of FCs associated with MDD (MDD-HC contrast) may not be comprehensive, as age-interacted FC associations may have been missed. Nonetheless, the prediction of RRS-B was made only for MDD, and age differences between groups did not affect this prediction. Another limitation is that we focused on the rumination portion of the RNT in MDD, neglecting the association with worry, another form of RNT that has been extensively discussed in anxiety disorders. Thus, the current FC findings concerning trait RNT are limited to rumination aspects in MDD. Additionally, it is important to acknowledge the limitation of the sample size in the current study. Despite implementing rigorous statistical evaluations, such as cross-validation and permutation tests, and employing multivariate approaches that can partially overcome the limitations of effect size, our sample may not be fully representative of the heterogeneous nature of the depressed population. Further investigations with larger samples are needed to draw more definitive conclusions about the association between RNT and RSFC and NTFC.

In conclusion, the results of the present study challenge the assumption that the resting state is equivalent to the negative thinking state in individuals with depression. While the resting state is often considered a proxy for the ruminative state, the results of this study suggest that resting state and negative thinking are not synonymous in depressed individuals. It is important to recognize that the functional implications of the resting state cannot be fully understood on the basis of resting state results alone. The value of resting state studies in depression should not be discounted; however, it is crucial to consider that interpretation of their functional implications requires additional information. Resting state, as an experimental procedure, does not necessarily reflect intrinsic functional activations (Finn, 2021), and the functional implications of resting state cannot be clarified without obtaining participants’ introspective reports (Gonzalez-Castillo et al., 2021). While the resting state may indicate abnormal brain activity in MDD, it may not fully capture the complexity of the rumination process. Negative thinking in depression involves dynamic interactions across multiple functional networks rather than being restricted to a specific brain network, which is not represented in the resting state.

## Acknowledgments

This work has been supported by The Laureate Institute for Brain Research and the National Institute of General Medical Sciences Center Grant Award Number (1P20GM121312). The content is solely the responsibility of the authors and does not necessarily represent the official views of the National Institutes of Health.

## Author contributions

Conceptualization, M.M. and A.T.; Methodology, M.M.; Investigation, M.M. and A.T.; Writing – Original Draft, M.M.; Writing – Review & Editing, M.M., A.T., S.M.G, M.L.R and M.P.P.

