## Supplementary Materials for "Trait repetitive negative thinking in depression is associated with functional connectivity in negative thinking state rather than resting state"

**Table S1**. Participants demographics and symptom scores.

| Group | N | Female | Age (year) | RRS total | RRS brooding | RRS depression | RRS reflection | MADRS |
|---|---|---|---|---|---|---|---|---|
| **HC** | 26 | 20 | 23.1±2.0 | 28.0±6.4 | 6.4±1.5 | 15.2±3.8 | 6.4±2.6 | N/A |
| **MDD** | 36 | 28 | 34.4±11.0 | 54.7±11.9 | 12.6±3.2 | 30.8±6.9 | 11.1±3.4 | 20.1±6.4 |
| **MDD-HC t-value** | | | 5.0*** | 10.3*** | 9.2*** | 10.4*** | 5.9*** | |

*** indicates $p < 0.001$, HC: Healthy controls, MDD: Major depressive Disorder, RRS: Ruminative Response Scale, MADRS: Montgomery-Asberg Depression Rating Scale

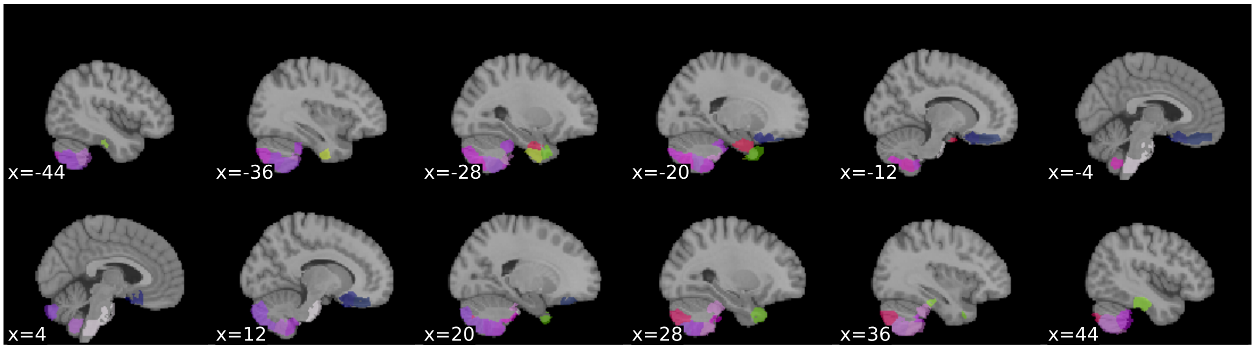

**Figure S1**. Regions excluded from CPM analysis in the Shen 268-node atlas due to limited coverage in the fMRI image. The atlas image was downloaded from https://www.nitrc.org/frs/download.php/7976/shen_1mm_268_parcellation.nii.gz
The indices of the excluded regions were 2, 4, 51, 55, 59, 100, 104, 107, 108, 109, 111, 112, 115, 116, 118, 119, 129, 130, 131, 135, 136, 137, 189, 196, 202, 235, 239, 240, 242, 243, 245, 246, 249, 250, 252, 256, 266, and 268.

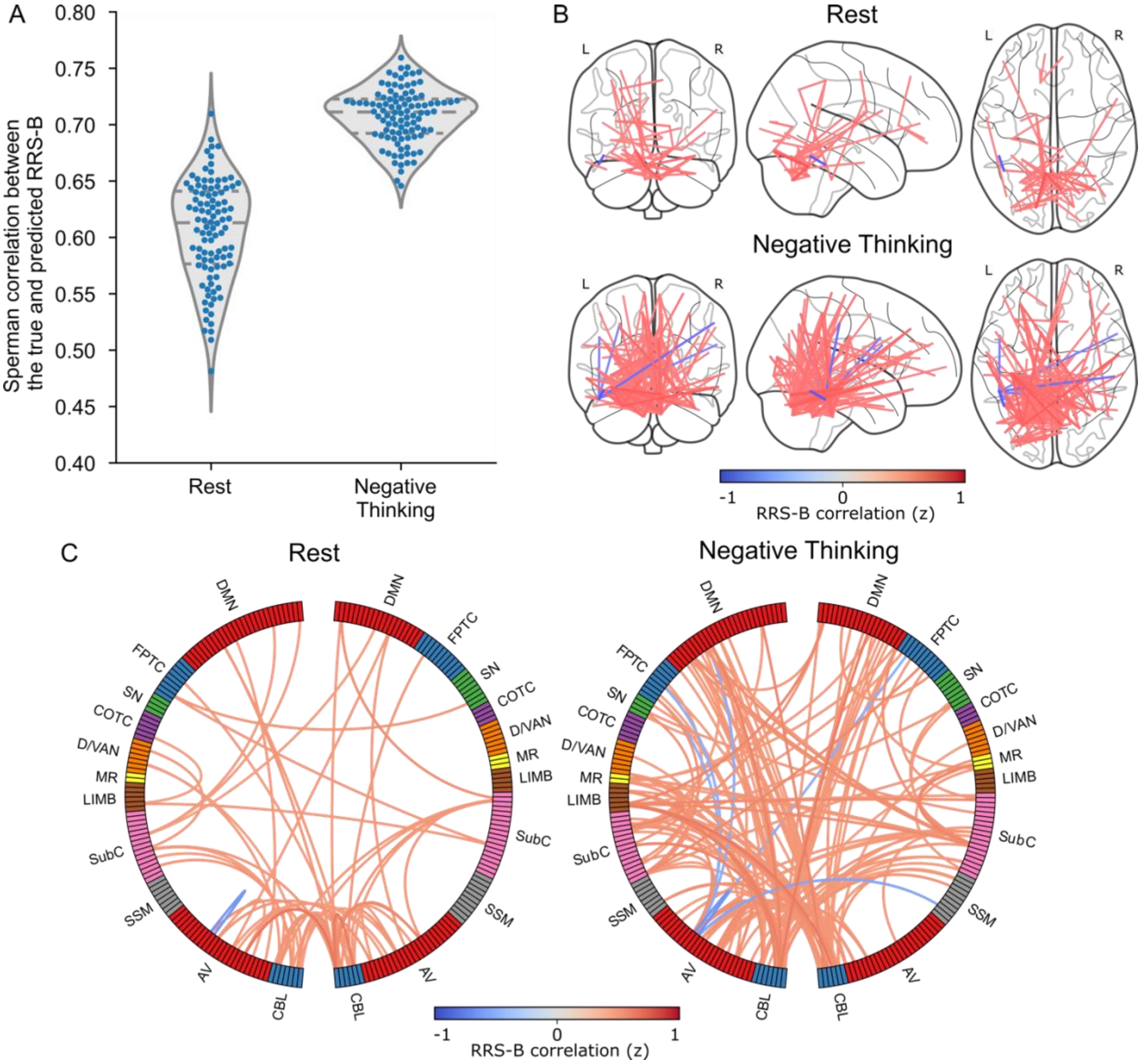

**Figure S2**. A. Distributions of the CPM prediction performance for individual RRS-B scores, including both HC and MDD groups. Each point indicates one iteration of the 5-fold cross-validation result (100 iterations with different random splits were performed). The violin plot and horizontal lines indicate the distribution curve and quartile positions. The median Spearman correlations between the true and predicted RRS-B scores were 0.613 ($p < 0.001$) for the resting state and 0.711 ($p < 0.001$) for the NT state. The difference in performance between the resting and NT states was not significant ($p = 0.247$). B. Plots of the connectivity selected by the CPM model. Connectivities selected by more than 85% cross-validation iterations were plotted on the glass brain. Line color indicates the connectivity correlation with RRS-B (z-transformed). C. Circle plots of the same connectivities as in B, summarized for each network region. Network labels are taken from Drysdale et al. (2017). These plots (B, C) show that the prediction performance was confounded by the significant group difference in the RRS-B score (SI Table S1), as the connectivity included in the prediction model overlapped with those informative for group classification (i.e., connectivity in the visual cortex and cerebellum regions). DMN: Default Mode Network, FPTC: Fronto-Parietal Task Control, SN: Salience Network, COTC: Cingulo-Opperculum Task Control, D/VAN: Doral Visual Attention Network, MR: Memory Retrieval, LIMB: default mode/limbic, SubC: Subcortical, SSM: Sensory SomatoMotor, AV: Auditory-Visual, CBL: Cerebellum.

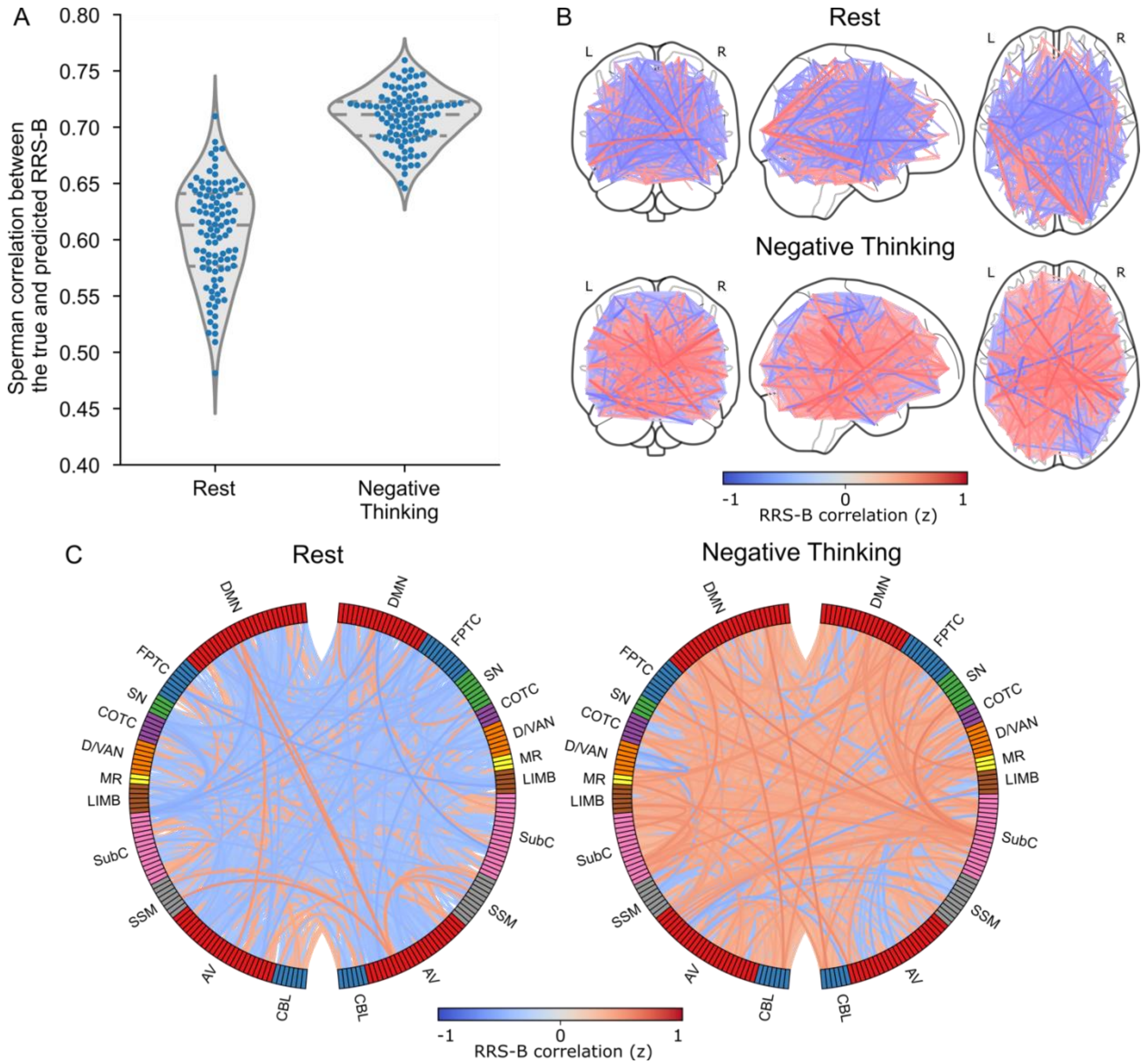

**Figure S3**. Same plot as Figure 2 in the main text, except that the connectivities selected by more than 50% cross-validation iterations are plotted in B and C.

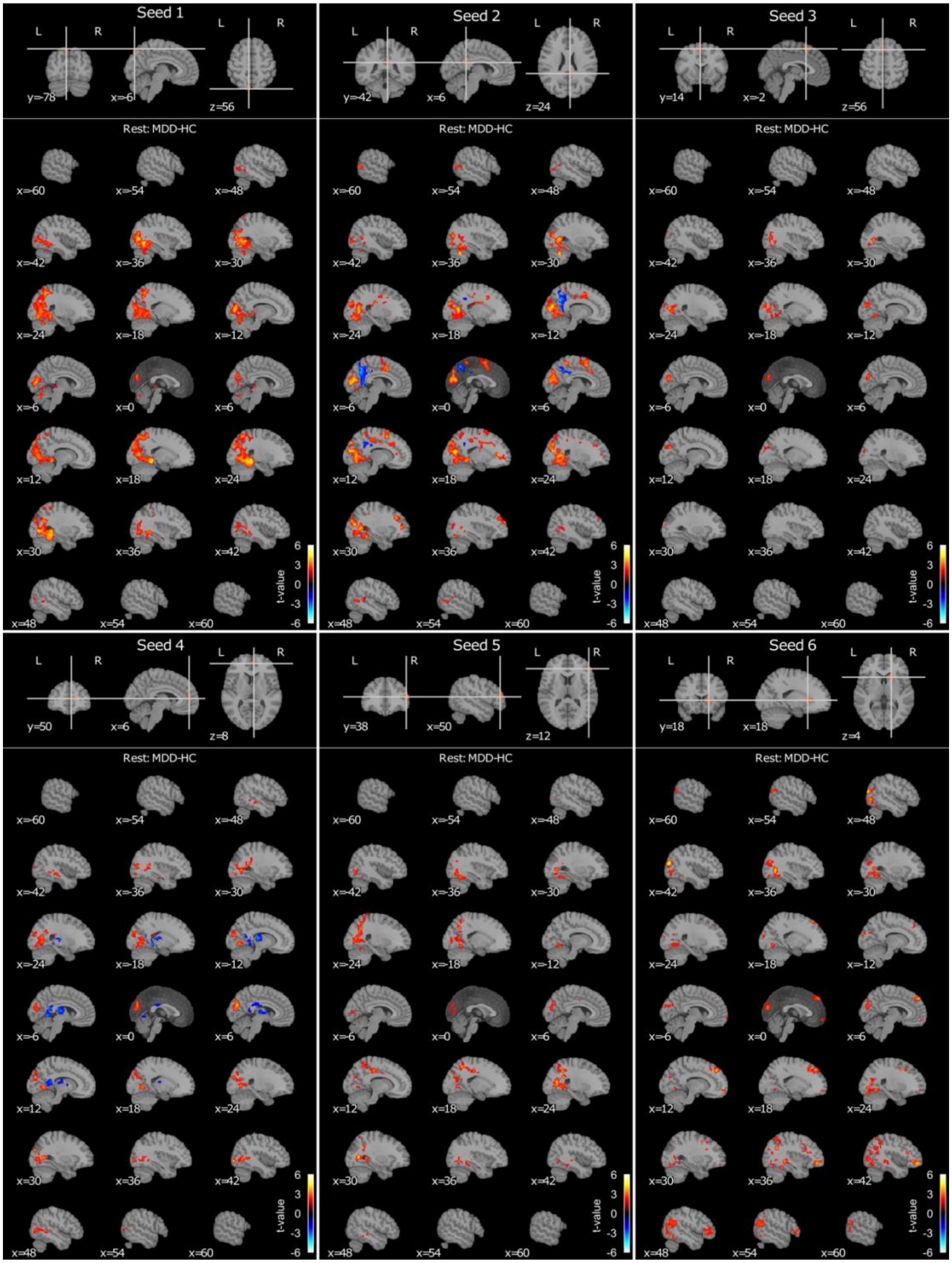

**Figure S4-1**. MDD-HC contrast connectivity maps in the MDMR post-hoc analysis for seeds with the significant contrast effect in the resting state. The seed index corresponds to Table 1. The map shows the t-value for MDD-HC contrast.

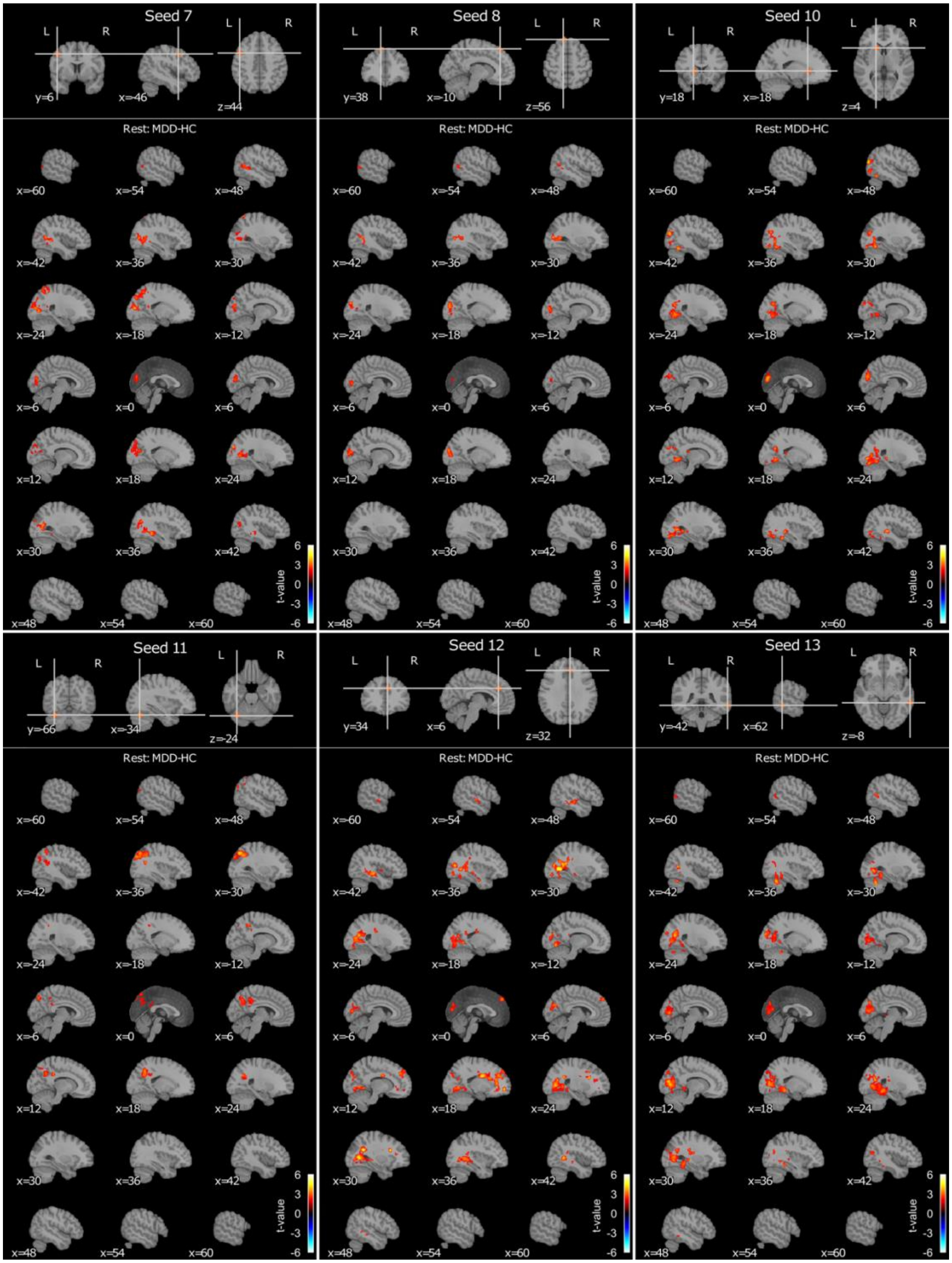

**Figure S4-2**. Continued from Figure S4-1.

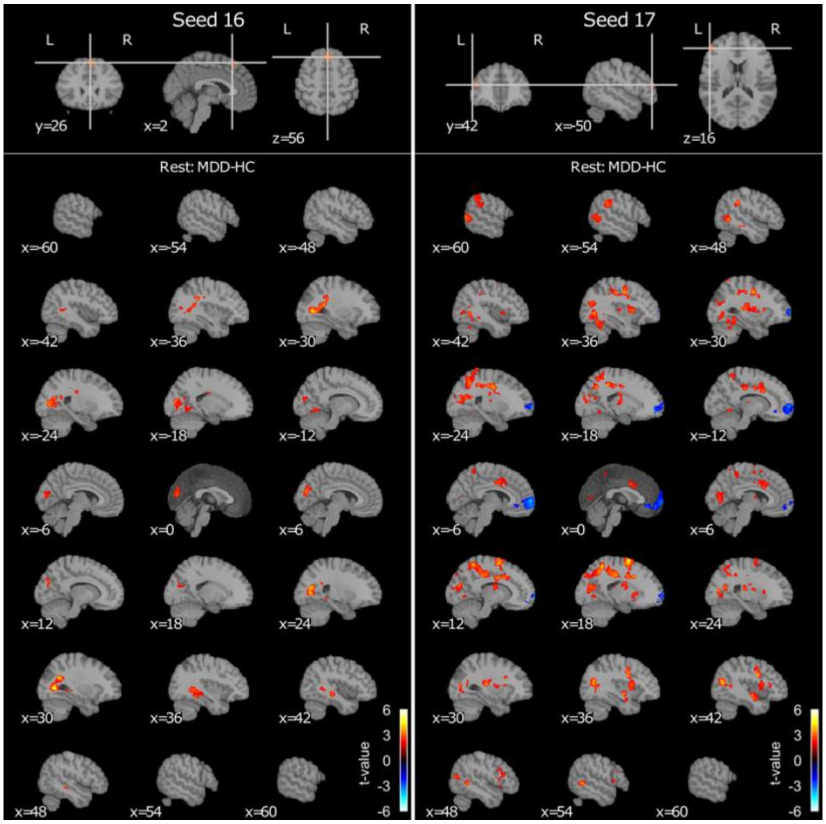

**Figure S4-3**. Continued from Figure S4-2.

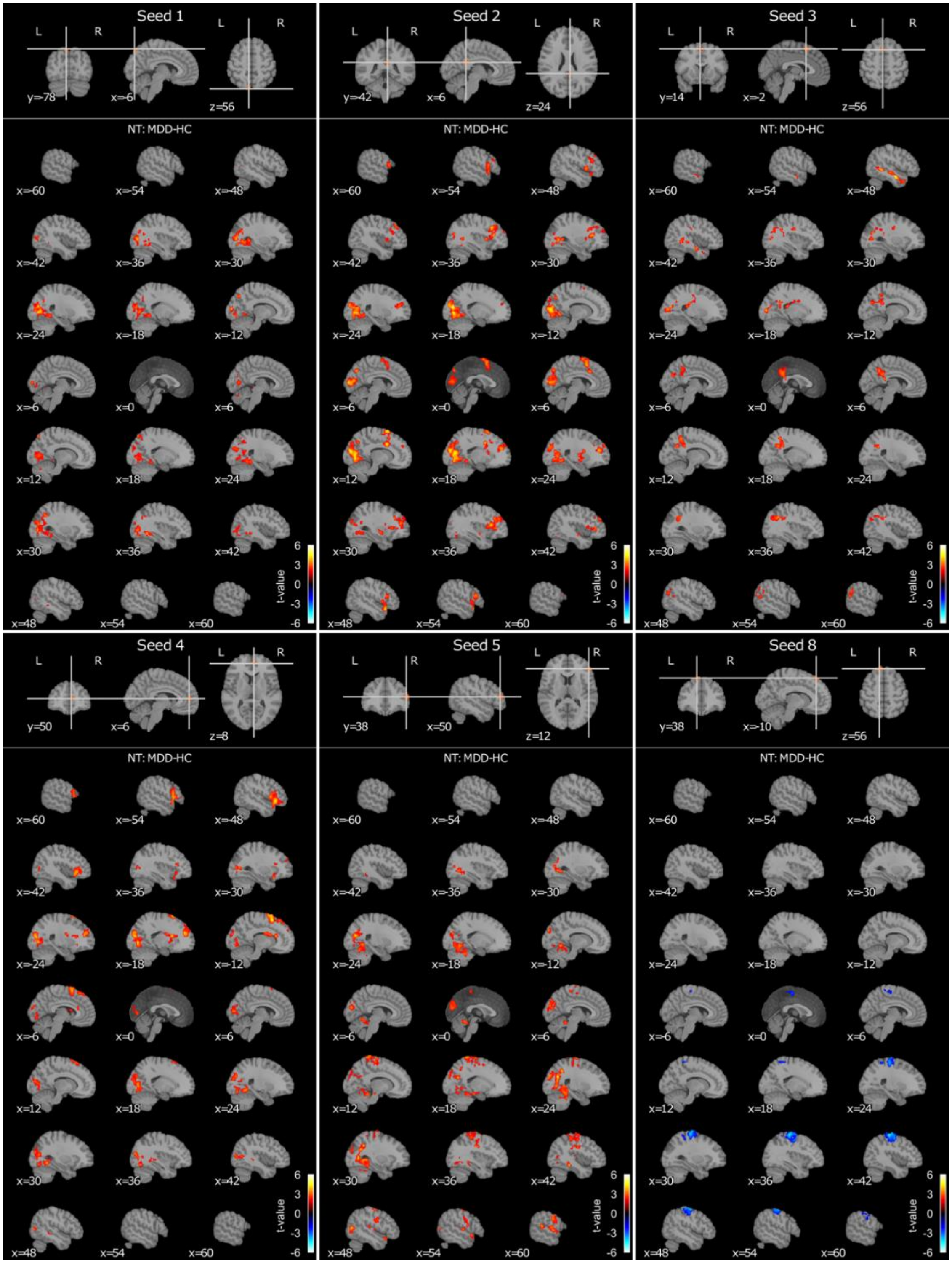

**Figure S5-1**. MDD-HC contrast connectivity maps in the MDMR post-hoc analysis for seeds with the significant contrast effect in the negative thinking (NT) state. The seed index corresponds to Table 1. The map shows the t-value for the MDD-HC contrast.

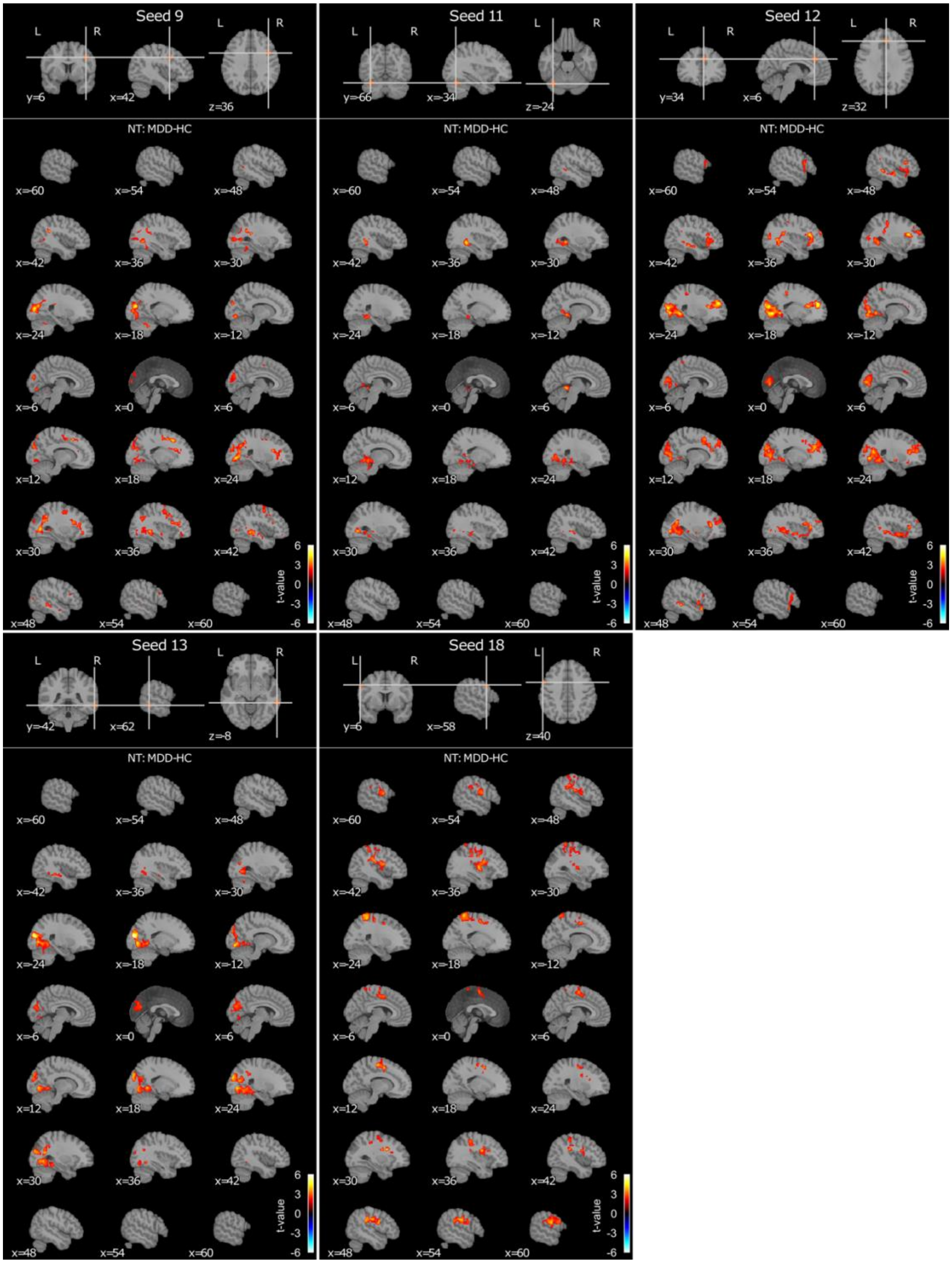

**Figure S5-2**. Continued from Figure S5-1.

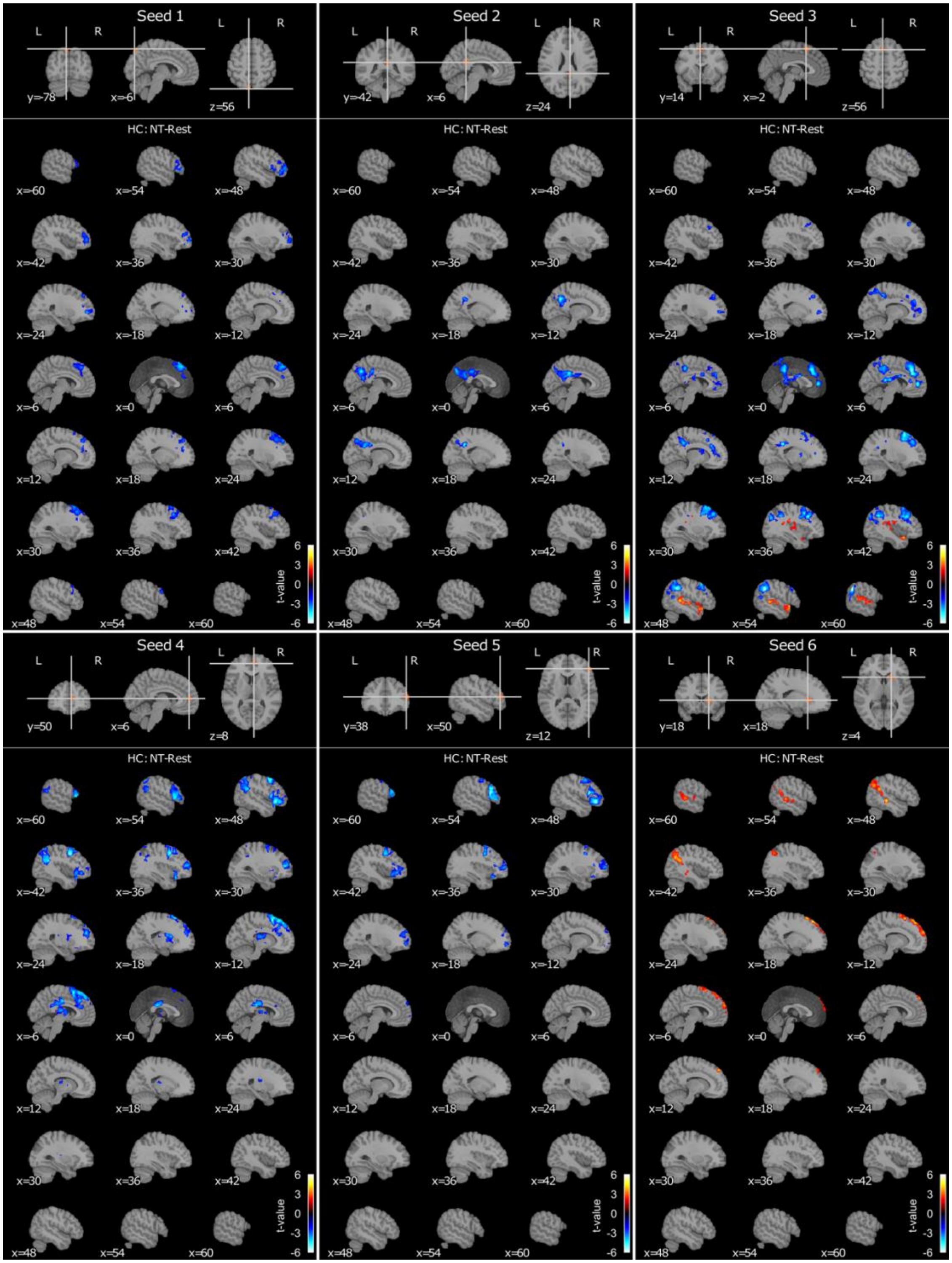

**Figure S6-1**. Negative thinking (NT)-Rest contrast connectivity maps in the MDMR post-hoc analysis for seeds with the significant contrast effect in the HC group. The seed index corresponds to Table 1. The map shows the t-value for the NT-rest contrast.

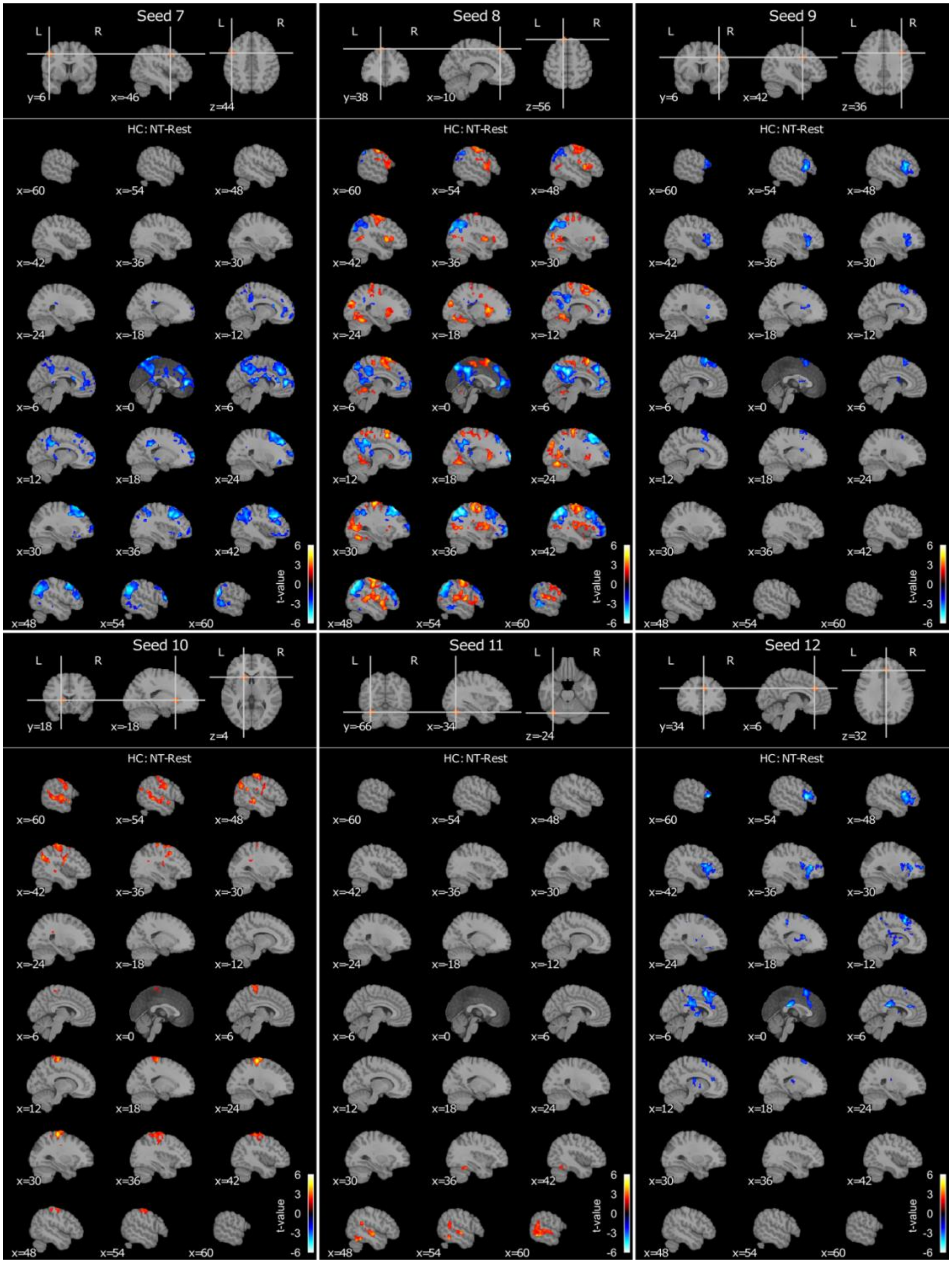

**Figure S6-2**. Continued form Figure S6-1.

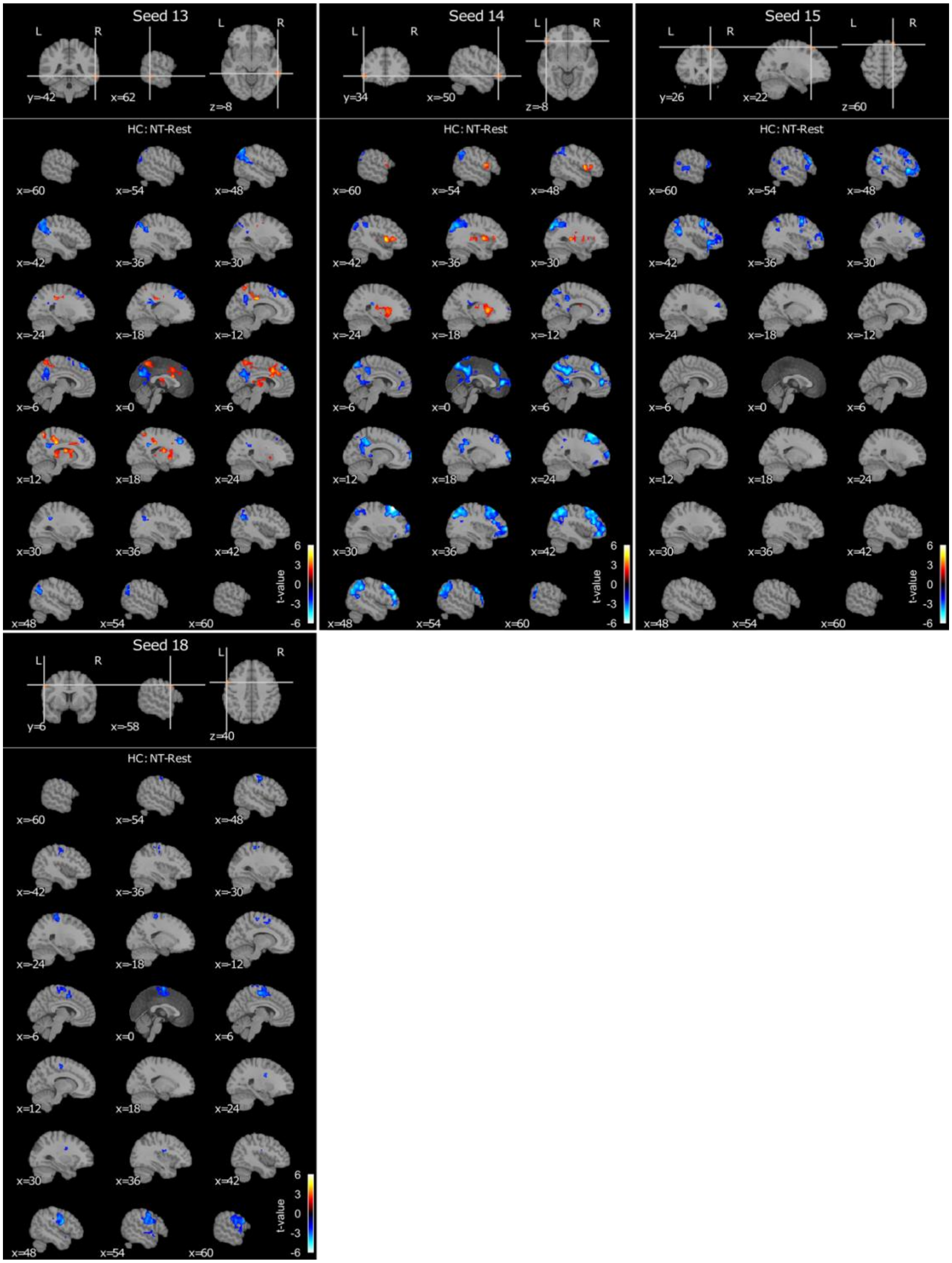

**Figure S6-3**. Continued from Figure S6-2.

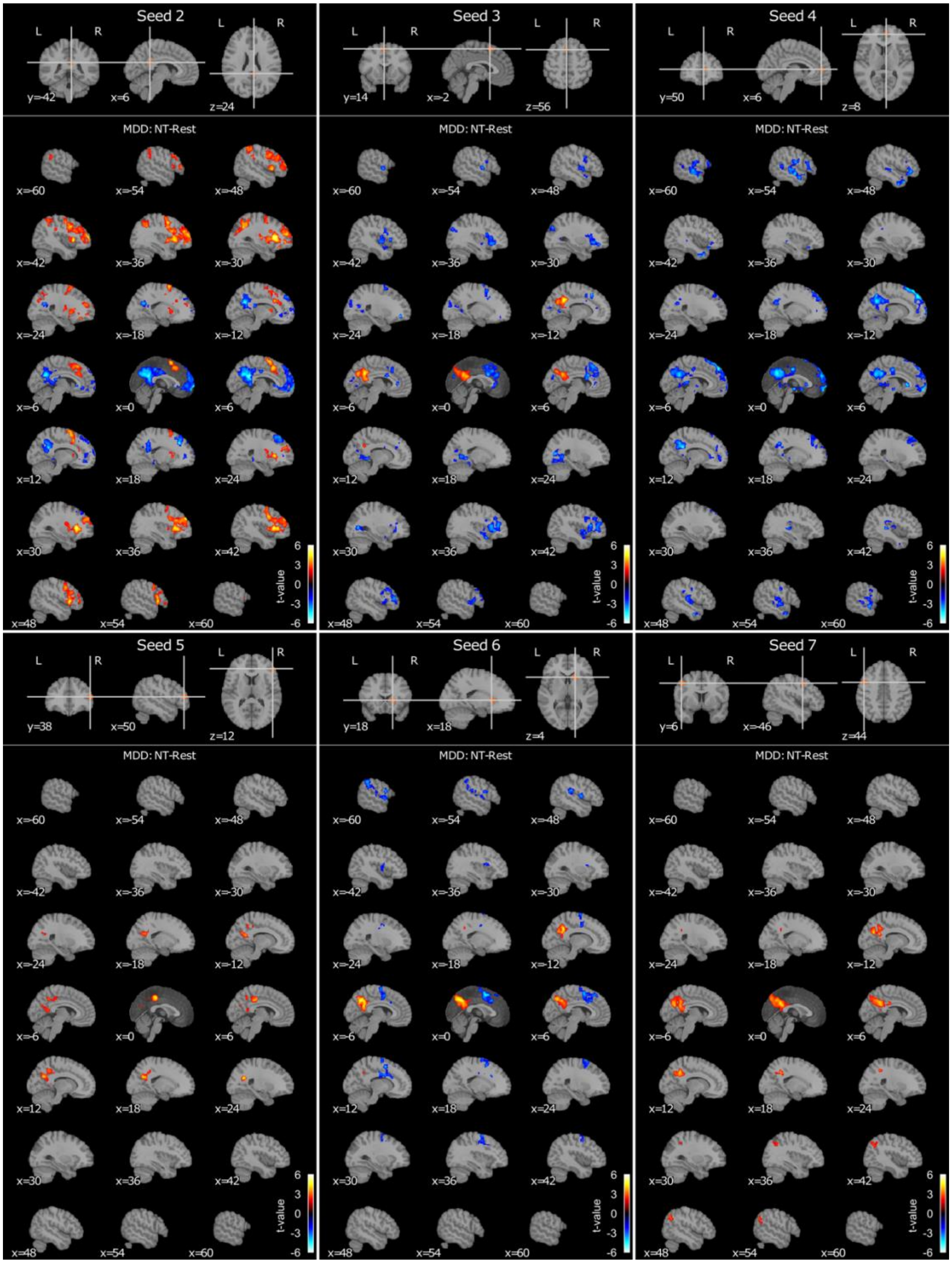

**Figure S7-1**. Negative thinking (NT)-Rest contrast connectivity maps in the MDMR post hoc analysis for seeds with the significant contrast effect in the MDD group. The seed index corresponds to Table 1. The map shows the t-value for the NT-rest contrast.

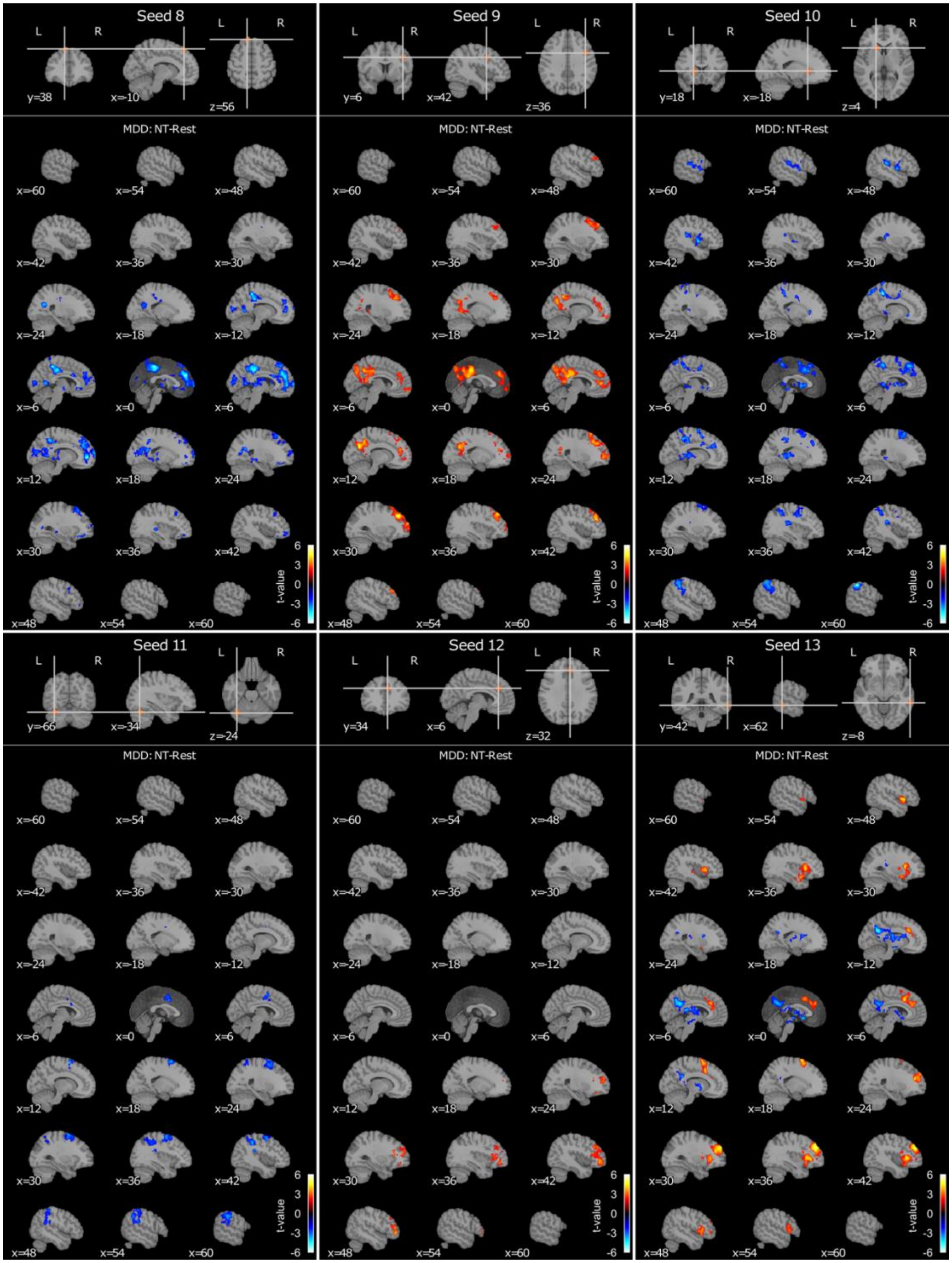

**Figure S7-2**. Continued from Figure S7-1.

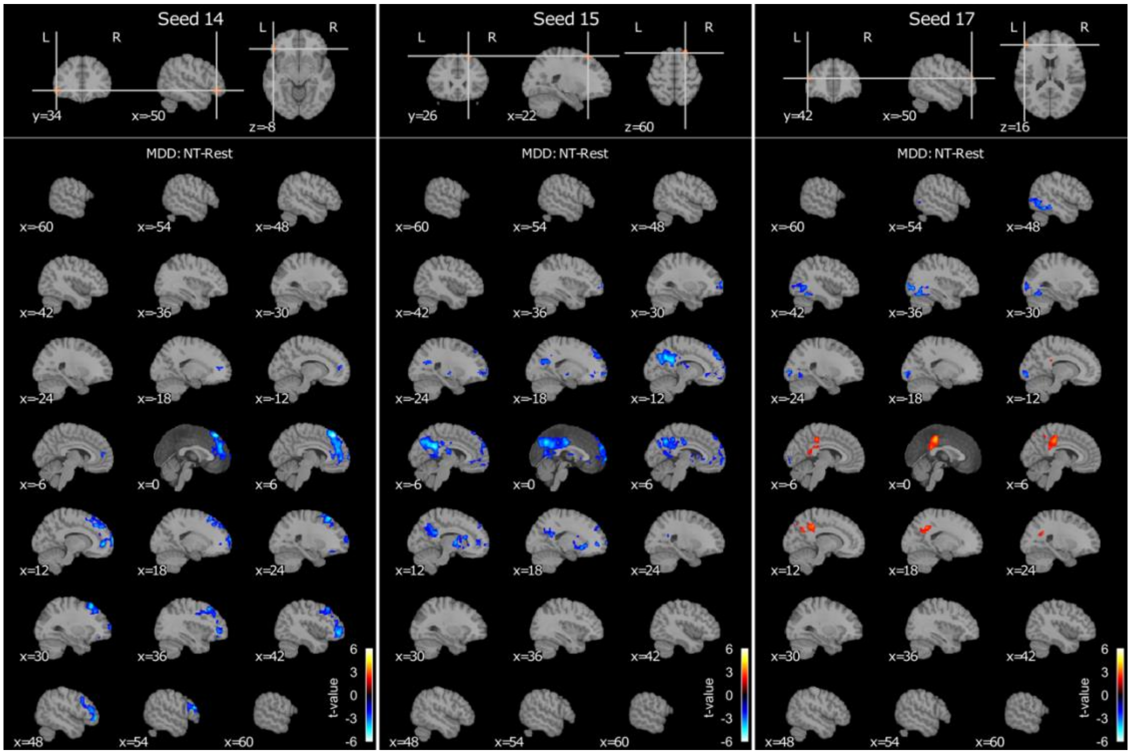

**Figure S7-3**. Continued from Figure S7-2.

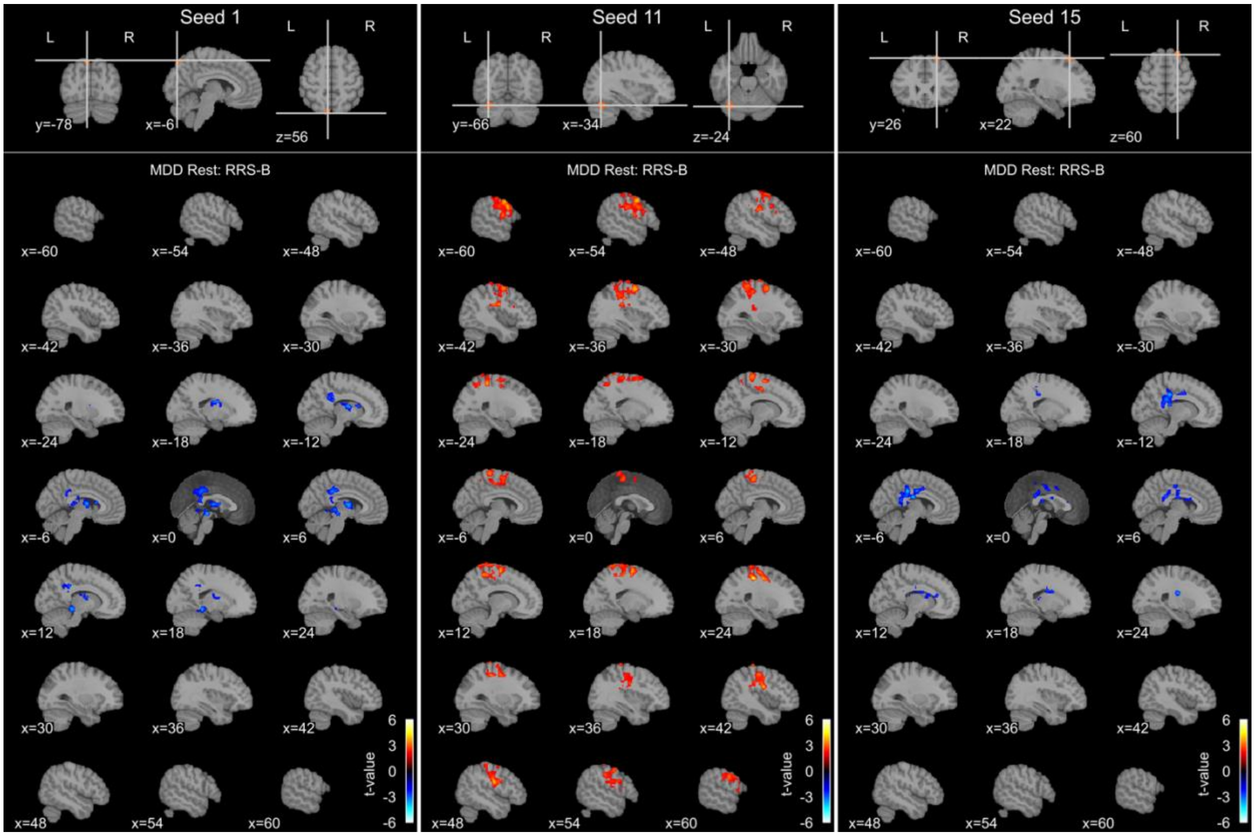

**Figure S8**. RRS-B association connectivity maps in the MDMR post-hoc analysis for seeds with the significant RRS-B effect in the resting state for the MDD group. The seed index corresponds to Table 1. The map shows the t-value for the RRS-B association.

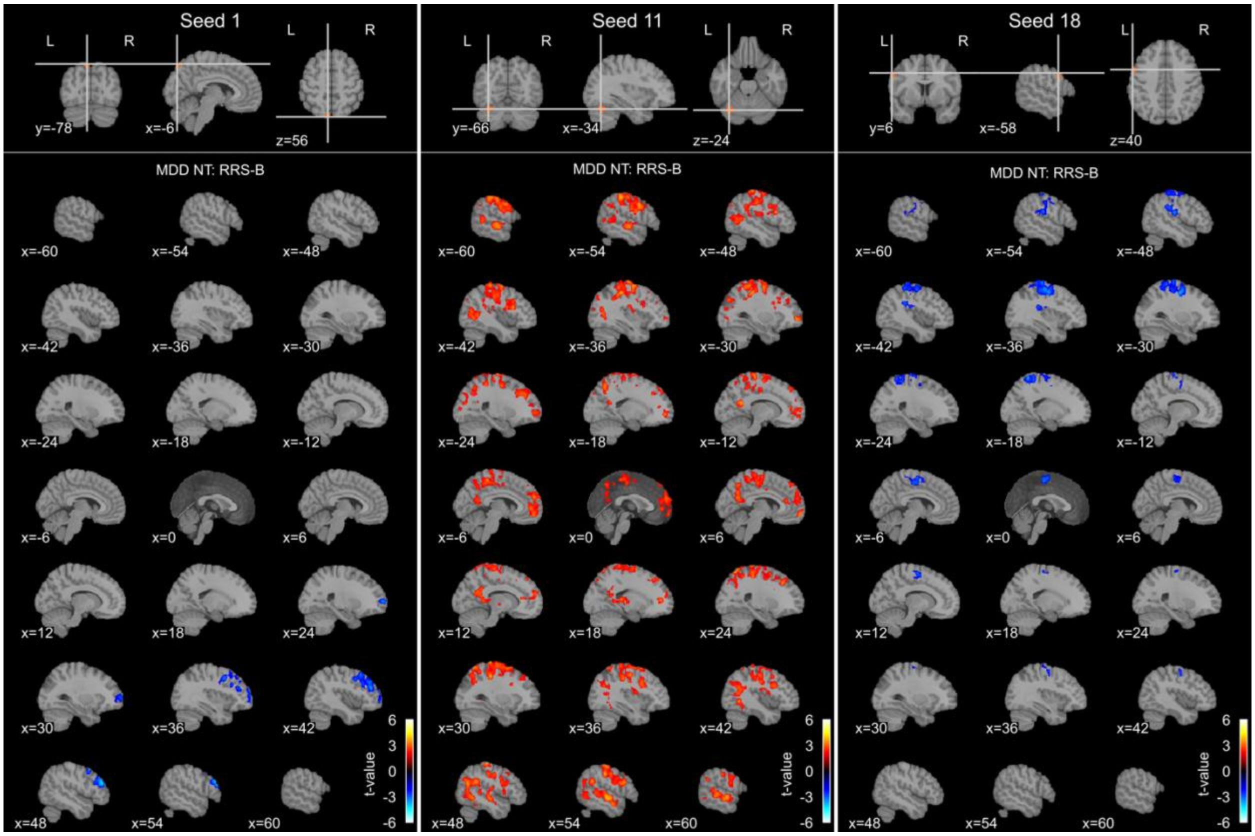

**Figure S9**. RRS-B association connectivity maps in the MDMR post-hoc analysis for seeds with the significant RRS-B effect in the negative thinking (NT) state for the MDD group. The seed index corresponds to Table 1. The map shows the t-value for the RRS-B association.
